# Syntenin and CD63 Promote Exosome Biogenesis from the Plasma Membrane by Blocking Cargo Endocytosis

**DOI:** 10.1101/2023.05.26.542409

**Authors:** Yiwei Ai, Chenxu Guo, Marta Garcia-Contreras, Laura S. Sánchez B., Andras Saftics, Oluwapelumi Shodubi, Shankar Raghunandan, Junhao Xu, Shang Jui Tsai, Yi Dong, Rong Li, Tijana Jovanovic-Talisman, Stephen J. Gould

## Abstract

Exosomes are small extracellular vesicles important in health and disease. Syntenin is thought to drive the biogenesis of CD63 exosomes by recruiting Alix and the ESCRT machinery to endosomes, initiating an endosome-mediated pathway of exosome biogenesis. Contrary to this model, we show here that syntenin drives the biogenesis of CD63 exosomes by blocking CD63 endocytosis, thereby allowing CD63 to accumulate at the plasma membrane, the primary site of exosome biogenesis. Consistent with these results, we find that inhibitors of endocytosis induce the exosomal secretion of CD63, that endocytosis inhibits the vesicular secretion of exosome cargo proteins, and that high-level expression of CD63 itself also inhibits endocytosis. These and other results indicate that exosomes bud primarily from the plasma membrane, that endocytosis inhibits their loading into exosomes, that syntenin and CD63 are expression-dependent regulators of exosome biogenesis, and that syntenin drives the biogenesis of CD63 exosomes even in Alix knockout cells.

## Introduction

Exosomes are ∼30-150 nm in diameter, have the same topology as the cell, and are highly enriched in specific exosomal proteins, especially the exosomal tetraspanins (e.g. CD81, CD9, and CD63 (*1-4*)) and their associated scaffolds (e.g. syntenin(*2, 5*)). Exosomes are released by all eukaryotic cells, and animals use exosomes for a wide variety of biological processes, including the transmission of signals and molecules to neighboring cells, modification of biofluids and extracellular structures, and protein quality control within exosome-producing cells(*6*). A clear understanding of exosome biogenesis is therefore critical to our understanding of basic cell biology, human health and disease, and the emerging field of exosome-based therapeutics(*7-10*). Human cells also release other types of extracellular vesicles (EVs), both small and large, yet exosomes are unique in their selective enrichment of specific cargo proteins, especially the exosomal tetraspanins.

Of the highly-enriched cargoes found in exosomes, CD63 is thought to be the cargo protein that best defines the exosome biogenesis pathway(*11-14*). It is widely presumed that this pathway requires the endocytosis and endosomal accumulation of CD63, where it recruits the cD63-binding protein syntenin, the syntenin-binding protein Alix, and the Alix-binding *endosomal sorting complexes required for transport* (ESCRT), presumably to drive the biogenesis of intralumenal vesicles (ILVs) that can later be secreted via endolysosomal exocytosis(*12, 15-23*). In support of this model, CD63 has been shown to binding syntenin directly, with the C-terminal four amino acids of CD63 being bound directly by syntenin’s PDZ domains and C-terminal peptide (syntenin amino acids 100-29), leaving syntenin’s N-terminal 100 amino acids free to bind Alix, through its three Alix-binding YPLxL motifs(*24*). These protein-protein interactions provide critical insights into the assembly of endosome-localized CD63/syntenin/Alix complexes, but its currently unclear whether these interactions are involved in the exosomal secretion of CD63, the lysosomal trafficking and degradation of CD63 and its partner proteins(*25-30*), or some other CD63-mediated process.

Contrary to this endosome-dependent model of exosome biogenesis, our group has established that the most highly enriched exosome cargo proteins (e.g. CD81, CD9, etc.) all reside at the plasma membrane(*3, 4, 31-33*), that targeting these plasma membrane-localized exosome cargoes to endosomes greatly reduces their exosomal secretion(*3, 4, 33*), and that that redirecting CD63 from endosomes to the plasma membrane greatly increased its exosomal secretion(*3, 4*). These results support an alternative hypothesis in which exosome biogenesis is mediated by a shared, stochastic mechanism that operates along the spectrum of plasma and endosome membranes, with most exosomes arising by direct budding from the cell surface(*3, 4, 6*).

One approach to testing between these models is to knockout these genes and ask whether loss of CD63, Alix, or syntenin causes a defect in exosome biogenesis. Although knockout of the CD63 or Alix genes failed to cause a defect in exosome biogenesis(*3*), knockout of syntenin caused an ∼50% reduction in the exosomal secretion of CD63(*13*), providing a genetic context for studying syntenin’s role in exosome biogenesis. Using a combination of genetic and cell biological studies, we show that syntenin plays a key role in the exosomal secretion of CD63 by blocking CD63 endocytosis, which allows it to accumulate at the primary site of exosome biogenesis, the plasma membrane. We also show that endocytosis is a general inhibitor of exosome cargo protein budding, and that high-level expression of CD63 induces its own exosomal secretion by saturating the clathrin adaptor AP-2 and blocking its endocytosis and the endocytosis of other lysosome membrane proteins.

## Results

### Syntenin expression induces the exosomal secretion of CD63

As noted above, knockout or silencing of the syntenin gene causes a selective defect in the exosomal secretion of CD63(*13, 21*). To determine whether loss of syntenin results in the same phenotype in 293F cells, we knocked out the syntenin gene (SDCBP) in this human cell line (***fig. S1***). The resulting F/SDCBP^-/-^ cell line displayed the same, CD63-selective phenotype as previously reported(*13, 21*), reducing the exosomal secretion of CD63 but showing no effect on the exosomal secretion of CD81 or CD9 (***fig. S1***).

To shed more light on syntenin’s role in exosome biogenesis, we asked whether its high-level expression was sufficient to induce the complementary phenotype of increased CD63 budding, and if so, whether this effect was selective to CD63. Towards this end, we created Tet-on 293F (FtetZ) cell lines that carry doxycycline-inducible, TRE3G-driven transgenes encoding (***i***) syntenin; (***ii***) the syntenin mutant ΔN100syntenin, which binds CD63 but not Alix; or (***iii***) the syntenin mutant synteninΔC23, which binds Alix but not CD63(*24*). These and control (FtetZ) cells were grown in the absence or presence of doxycycline, followed by collection of cell and exosome fractions, measurement of transgene-encoded syntenin expression by qRT-PCR (***fig. S2***), and interrogation of cell and exosome samples by immunoblot to measure the exosomal secretion of CD63, CD81, and CD9 (***Fig. 1A-C***).

**Figure 1.**
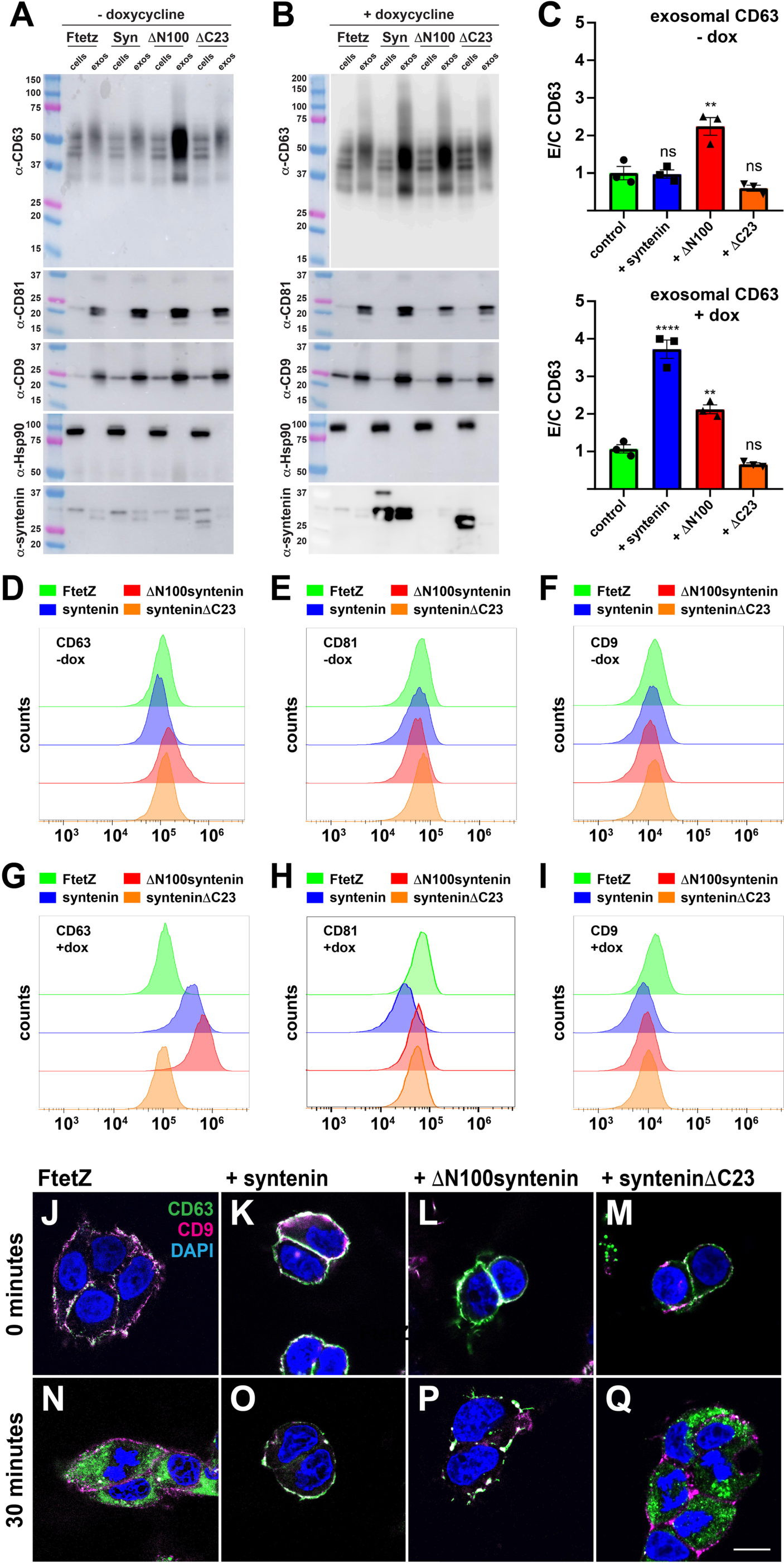
Syntenin induces the exosomal secretion of CD63 by blocking CD63 endocytosis. Immunoblot analysis of cell and exosome lysates collected from FtetZ and FtetZ cells carrying TRE3G-regulated transgenes encoding (Syn) WT human syntenin, (N100) N100syntenin, or (C23) syntenin C23, grown in the (**A**) absence or (**B**) presence of doxycycline. Immunoblots were probed using antibodies specific for the exosome cargo proteins CD63, CD9, and CD81, the cytoplasmic protein Hsp90, and syntenin. The anti-syntenin antibody was raised against an N-terminal fragment and does not bind N100syntenin. Similar results were observed in multiple trials (n = 4). (**C**) Bar graphs showing the mean exosome/cell ratio of CD63, +/- standard error of the mean (s.e.m.) in the experiments of A and B. Statistical significance was assessed by ANOVA, with ** denoting *p* value <0.01 and **** denoting a *p* value <0.0001. (**D-I**) Flow cytometry histograms of FtetZ and FtetZ cells carrying TRE3G-regulated transgenes encoding (Syn) WT human syntenin, (N100) N100syntenin, or (C23) syntenin C23, grown in the (D-F) absence or (G-I) presence of doxycycline, chilled and stained for the cell surface abundance of (D, G) CD63, (E, H) CD81, or (F, I) CD9. (**J-Q**) Confocal fluorescence microscopy of (J, N) FtetZ cells, and FtetZ cells carrying TRE3G-regulated transgenes encoding (K, O) syntenin, (L, P) N100syntenin, or (M, Q) syntenin C23 that had been chilled, incubated with fluorescently tagged antibodies specific for (green) CD63 and (pink) CD9, then warmed and then fixed and imaged after (J-M) 0 minutes at 37°C or (N-Q) 30 minutes at 37°C. Bar, 10 um. Data is from three independent trails.

In the absence of doxycycline, all three syntenin transgene-encoded mRNAs were expressed at similar levels (***fig. S2***). This baseline expression was quite low, as we could not detect the syntenin protein by immunoblot in either FtetZ cells or FtetZ::syntenin cells (***Fig. 1A***). Nevertheless, the low baseline expression of ΔN100syntenin triggered a significant increase in the exosomal secretion of CD63, indicating that this N-terminal syntenin truncation mutant was a particularly potent inducer of CD63 exosome biogenesis, even though it no longer contained syntenin’s Alix-binding sites. Furthermore, when these same four cell lines were grown in the presence of doxycycline, we found that high-level expression of the syntenin and ΔN100syntenin transgenes both induced the exosomal secretion of CD63, whereas synteninΔC23 did not (***Fig. 1B, C***). Furthermore, these effects were selective to CD63, as high-level expression of syntenin or ΔN100syntenin had no effect on the exosomal secretion of either CD81 or CD9 (***Fig. 1A-C***). Similar results were observed in mouse NIH3T3 fibroblasts (***fig. S3***).

### Syntenin induces the plasma membrane accumulation of CD63

These immunoblots confirmed our prior discovery that CD81 and CD9 are loaded into exosomes far more efficiently than CD63 (15-fold and 5-fold, respectively)(*1, 3, 4*), showed that high-level expression allows CD63 to bud from cells at efficiencies that rival CD81 and CD9, and in so doing raised the possibility that syntenin was inducing the exosomal secretion of CD63 by relieving an inhibition of CD63’s exosomal secretion. As for what this inhibitory effect might be, we previously discovered that endocytosis inhibits CD63’s exosomal secretion by ∼6-fold(*3, 4*). To determine whether syntenin might be inhibiting the endocytosis of CD63, we used flow cytometry to measure the plasma membrane abundance of CD63, CD9, and CD81 in uninduced and doxycycline-induced cultures of these same four cell lines. At baseline, plasma membrane CD63 was ∼80% higher in ΔN100syntenin cells and was unchanged in cells carrying the syntenin or synteninΔC23 transgenes (n = 3, *p* <0.0005), while all four cell lines had similar levels of cell surface CD81 and CD9 (***Fig. 1D-F***). As for the doxycycline-induced cells, high-level expression of syntenin or ΔN100syntenin increased the plasma membrane abundance of CD63 by 3-fold and 5-fold, respectively (n = 3, *p* <0.0005), whereas synteninΔC23 had no effect, and the plasma membrane levels of CD81 and CD9 were largely unchanged (***Fig. 1G-I***). Similar results were observed in Hela cells (***Fig. S3***).

### Syntenin blocks CD63 endocytosis

The preceding results predict that high level expression of syntenin or ΔN100syntenin will block the endocytosis of CD63, as syntenin binds directly to the same C-terminal four amino acids of CD63 that comprise its endocytosis signal, -YEVMcooh, which is also bound by the mu2 subunit of the clathrin adaptor AP-2(*24, 34, 35*). We therefore seeded these four cell lines onto glass coverslips, incubated them in doxycycline-containing media overnight, then shifted them to 4°C, stained them with fluorescently tagged antibodies to (green) CD63 and (pink) CD9 (also at 4°C), washed (also at 4°C), then either fixed them immediately (t = 0) or following incubation at 37°C for 30 minutes (t = 30). Examination of these cells by confocal fluorescence microscopy confirmed that the control FtetZ cells endocytosed CD63 from the cell surface, leading to the rapid accumulation of CD63-antibody complexes in internal compartments scattered throughout the cytoplasm, whereas CD9 remained at the cell surface (***Fig. 1J, N***). In contrast, cells expressing either WT syntenin (***Fig. 1K, O***) or ΔN100syntenin (***Fig. 1L, P***) failed to endocytose CD63, demonstrating that high-level expression of syntenin had indeed blocked the endocytosis of CD63. This phenotype was specific for CD63-binding forms of syntenin, as high-level expression of synteninΔC23 failed to inhibit the endocytosis of CD63 (***Fig. 1M, Q***). These results are consistent with those of Latysheva et al.(*24*), who previously established that syntenin is an expression-dependent inhibitor of CD63 endocytosis.

### Syntenin drives the loading of CD63 into plasma membrane-derived, CD81/CD9 exosomes

We next tested whether high-level expression of syntenin affects the overall yield of exosome-sized EVs by counting their concentration in each of the above cultures. Small EVs were collected from each culture supernatant (by clarifying filtration, concentrating filtration, and size exclusion chromatography) and then interrogated by nanoparticle tracking analysis (NTA). Exosome yields were similar among all four cell lines and were not altered by addition of doxycycline (***Fig. 2A***), indicating that high-level expression of syntenin induced the exosomal secretion of CD63 without increasing the production of exosome-sized vesicles.

**Figure 2.**
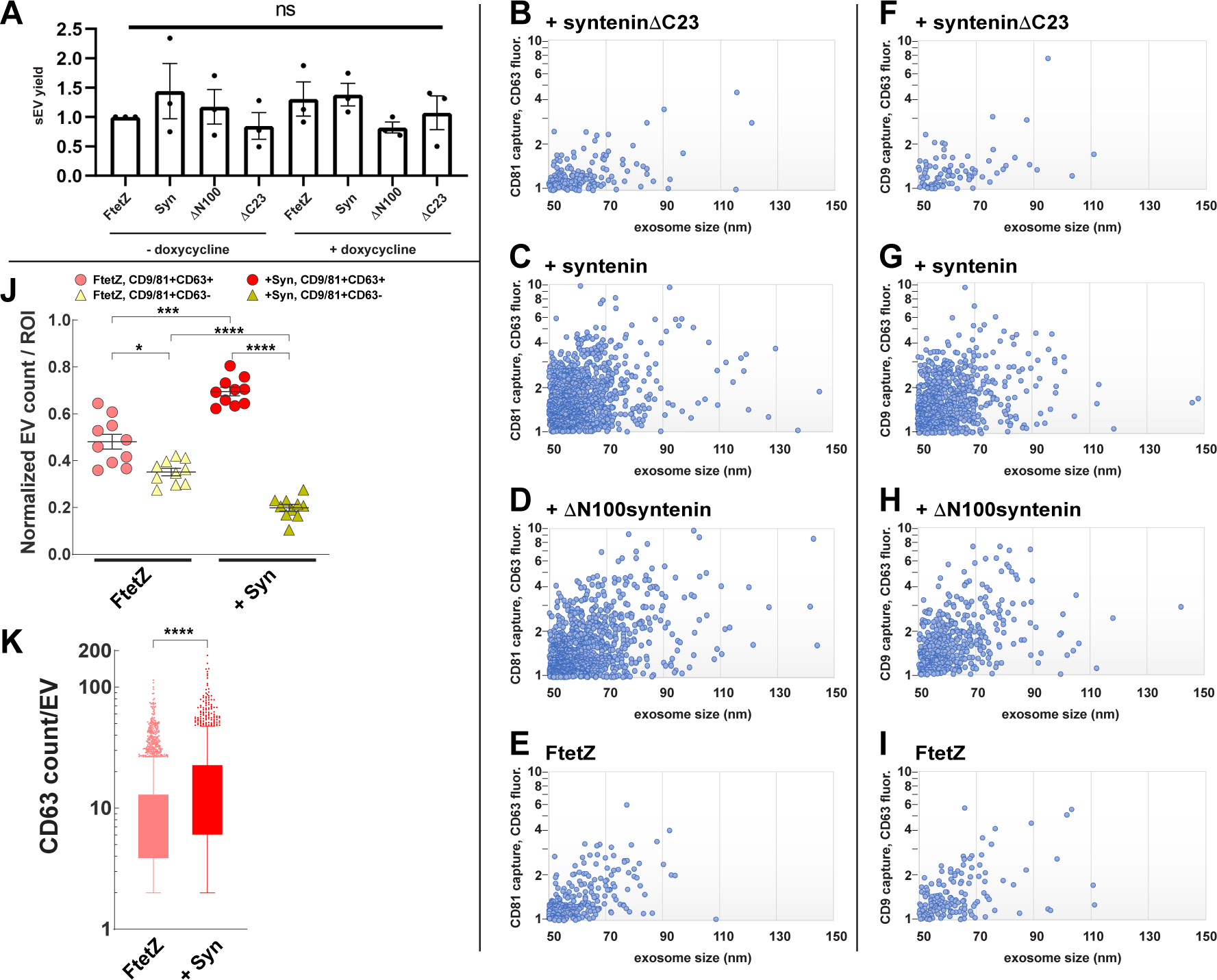
Syntenin drives the loading of CD63 into CD81 and CD9 exosomes. (**A**) Bar graph showing the yield of sEVs in exosome preparations collected from uninduced and induced FtetZ, FtetZ::syntenin. FtetZ:: N100syntenin, and FtetZ::syntenin C23 cells. Bar height represents the average, error bars represent the standard error of the mean, and ANOVA showed that none of the differences approached the cutoff for statistical significance (*p* <0.05). (**B-E**) Scatter plot of (y-axis) anti-CD63 fluorescence (arbitrary units) of CD81-containing exosomes, plotted versus (x-axis) exosome size, as determined by SPIR-IFM imaging of exosomes produced by (B) FtetZ::syntenin C23 cells, (C) FtetZ::syntenin cells, (D) FtetZ:: N100syntenin, and (E) FtetZ cells. (**F-I**) Scatter plot of (y-axis) anti-CD63 fluorescence (arbitrary units) of CD9-containing exosomes, plotted versus (x-axis) exosome size, as determined by SPIR-IFM imaging of exosomes produced by (F) FtetZ::syntenin C23 cells, (G) FtetZ::syntenin cells, (H) FtetZ:: N100syntenin, and (I) FtetZ cells. (**J, K**) qSMLM-determined exosome immunophenotypes of CD81/CD9-immunopurified exosomes collected from doxycycline-induced FtetZ and FTetZ::syntenin cells, stained with AF647-labeled anti-CD63 and CF568-labeled anti-CD81 and anti-CD9 antibodies. (J) Scatter dot plot displaying the proportion of CD81/CD9-positive exosomes produced by FtetZ and FtetZ::syntenin cells that were (red dots) CD63 positive or (gold triangles) CD63 negative. Data are from 10 independent regions of interest (ROI) for each sample from two coverslips. Wide bars denote the average, and error bars denote the s.e.m. ****, ***, and * denote *p* values of <0.00005, <0.0005, and <0.05 from Dunnett’s T3 multiple comparisons test. (K) Box and whisker plot showing the number of detected CD63 molecules per CD81/CD9-positive exosomes that had been produced by (left) FtetZ cells and (right) FtetZ::syntenin cells. **** denotes *p* value of <0.000001 from an unpaired t test with Welch’s correction. Data is from two independent trials.

We next tested whether syntenin induced the loading of CD63 into plasma membrane-derived exosomes, which are marked by CD81 and CD9(*3, 4, 11*). Towards this end, we interrogated the doxycycline-induced exosome fractions using the combined technologies of single-particle interferometric reflectance (SPIR) imaging, provides label-free measurement of individual exosome sizes, and conventional immunofluorescence microscopy (IFM), which allowed us to quantify the abundance of CD63 on each of thousands of individual exosomes(*3, 4, 36*). In brief, each exosome sample was subject to immunopurification on SPIR imaging chips that had been functionalized with a monoclonal antibody specific for human CD81, followed by staining with an Alexa Fluor 647 (AF647)-labeled antibody specific for CD63. Samples were then washed, fixed, and examined by SPIR-IFM imaging, allowing us to measure the sizes and CD63 levels on hundreds of individual CD81-positive exosomes produced by each cell line. CD63 fluorescence intensity was gated to exclude low and ambiguous signals, and the resulting plots of exosome size versus CD63 staining intensity revealed that CD63 was abundant on only ∼7% of CD81 exosomes produced by FtetZ::*TRE3G*-synteninΔC23 cells (152/2044), that expression of syntenin increased this to ∼40% (988/2389), and that expression of ΔN100syntenin increased to ∼35% (991/2791). As for the exosomes produced by FtetZ cells, ∼15% of CD81-positive exosomes carried this level of CD63 (250/1690), about twice the level seen in doxycycline-induced FtetZ::*TRE3G*-synteninΔC23 cells.

We also performed parallel studies in which the exosomes were immunopurified on SPIR imaging chips that had been functionalized with an anti-CD9 monoclonal antibody, then stained with AF647-labeled anti-CD63 antibodies. SPIR-IFM imaging of these exosomes revealed that high levels of CD63 were detected on ∼6% of CD9 exosomes produced by FtetZ::*TRE3G*-synteninΔC23 cells (85/1440), ∼35% of CD9 exosomes produced by FtetZ::*TRE3G*-syntenin cells (721/2059), ∼35% of CD9 exosomes produced by FtetZ::*TRE3G*-ΔN100syntenin cells (433/1267), and ∼16% of CD9 exosomes produced by FtetZ cells (159/966) (***Fig. 2F-I***).

As an orthologous approach to single exosome immunophenotyping, we interrogated exosomes from control and syntenin-expressing cells by quantitative single-molecule localization microscopy (qSMLM). In brief, CD81/CD9-positive exosomes were immunopurified on glass coverslips functionalized with a mixture of monoclonal antibodies specific for human CD81 and CD9, then stained with an AF647-labeled antibody specific for human CD63 and a pair of AF568-labeled antibodies specific for CD81 and CD9. The resulting samples were then washed, fixed, and examined by qSMLM. Of the CD81/CD9-positive exosomes produced by doxycycline-induced FtetZ and FtetZ::*TRE3G*-syntenin cells, we found that FtetZ exosomes contained an average of 10 detected CD63 molecules (coefficient of variation (c.v.) 102%) whereas FtetZ::syntenin exosomes contained an average of 18 detected CD63 molecules (c.v. 102%) (***Fig. 2J***). Also, when we calculated the percentage of CD81/CD9-positive exosomes that lacked detectable levels of CD63, we found that it fell from ∼35% for exosomes produced by FtetZ cells to ∼15% for exosomes produced by syntenin-expressing cells (***Fig. 2K***). Syntenin had no substantive effect on exosome size, as exosomes produced by FtetZ cells and FtetZ::*TRE3G*-syntenin cells had similar diameters (mean of 107nm (c.v. 26%) and 113 nm (c.v. 30%), respectively).

### Syntenin-induced exosome biogenesis is an Alix-independent process

Numerous studies have proposed that Alix plays a critical role in exosome biogenesis, and in particular, that Alix drives exosome biogenesis by linking CD63-syntenin complexes to the ESCRT machinery(*12, 15-23*). We therefore tested whether loss of Alix prevents the syntenin-induced exosomal secretion of CD63. Specifically, we created HtetZ/Alix^-/-^ cells, a Tet-On derivative of an Alix^-/-^ HEK293 cell line(*3*), then modified this HtetZ/Alix^-/-^ cell line to carry the doxycycline-regulated transgenes encoding syntenin, ΔN100syntenin, or synteninΔC23. We then examined uninduced and doxycycline-induced cultures of these cell lines by flow cytometry. Even though these Alix^-/-^ cells lack Alix protein, high-level expression of syntenin or ΔN100syntenin still induced the selective plasma membrane accumulation of CD63 (***fig. S4A-C***). In addition, immunoblot of cell and exosome fractions from these Alix^-/-^ cell lines demonstrated that high-level expression of syntenin or ΔN100syntenin still induced the exosomal secretion of CD63 (***fig. S4D, E***). Thus, Alix is not required for the syntenin-induced exosomal secretion of CD63, consistent with the fact that Alix knockout cells show no defect in the release of exosome-sized vesicles or the exosomal secretion of CD81, CD9, or CD63(*3*), and that ΔN100syntenin induced the exosomal secretion of CD63 (***Fig. 1A-C***), even though it lacks all three of syntenin’s Alix-binding YPLxL motifs(*24*).

### General inhibitors of endocytosis induce the exosomal secretion of CD63

The hypothesis that emerges from these data is that syntenin drives the exosomal secretion of CD63 indirectly, by blocking its endocytosis and inducing its accumulation at the primary site of exosome biogenesis, the plasma membrane. If this hypothesis is correct, then other, mechanistic distinct inhibitors of CD63 endocytosis should also induce the exosomal secretion of CD63. To test this prediction, we used Cas9 to generate mutations in the AP2M1 gene, which encodes the mu2 subunit of the clathrin adaptor AP-2(*35, 37-39*). This effort yielded a single 293F AP2M1^-/-^ cell line, and sequence analysis of these cells revealed that this cell line has a large deletion on AP2M1 allele #1 and a 1 codon deletion (Ile63) on AP2M1 allele #2 (***fig. S5***). To determine whether this F/AP2M1^-^ /- cell line was defective in CD63 endocytosis, we grew it and control 293F cells on glass coverslips, chilled them to 4°C, incubated them with fluorescently tagged antibodies specific for CD63 and CD9 (also at 4°C), then washed the cells and fixed them either immediately (t = 0) or after a 30 minutes-long incubation at 37°C (t = 30). Confocal immunofluorescence microscopy showed that 293F cells rapidly endocytosed CD63 from the cell surface (***Fig. 3A, B***) but that F/AP2M1^-/-^ cells did not (***Fig. 3C, D***), demonstrating that F/AP2M1^-/-^ cells were indeed defective in CD63 endocytosis. Not surprisingly, interrogation of these cells by flow cytometry revealed a selective, ∼4-fold increase in plasma membrane CD63 abundance, but no change in cell surface CD81 or CD9 (***Fig. 3E***). Importantly, immunoblot analysis of cell and exosome fractions from these cells showed that F/AP2M1^-/-^ cells display an ∼2-fold increase in their exosomal secretion of CD63 (***Fig. 3F, G***), demonstrating that a mechanistically distinct inhibition of CD63 endocytosis was sufficient to induce a selective increase in the exosomal secretion of CD63.

**Figure 3.**
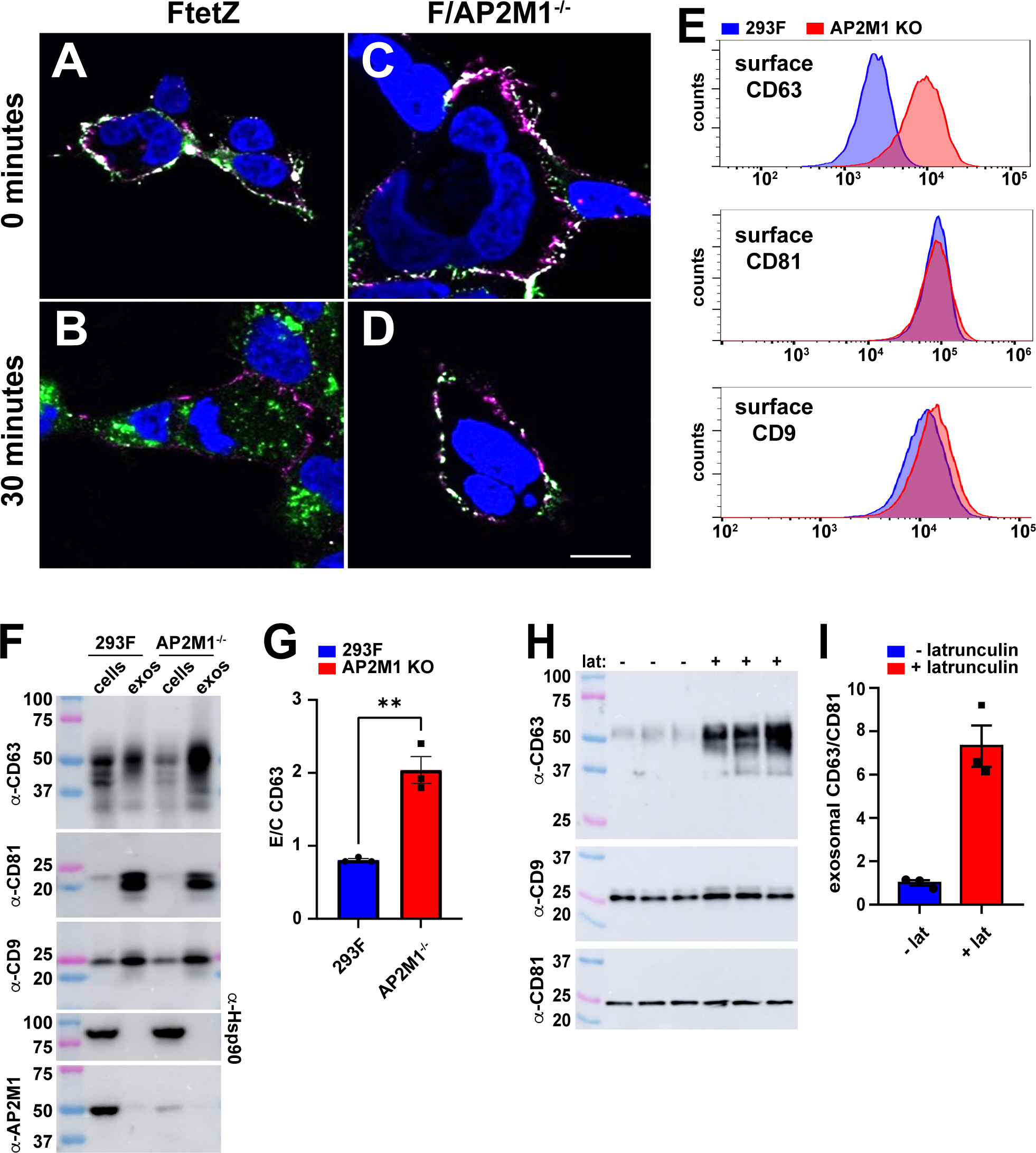
Inhibitors of endocytosis induce the exosomal secretion of CD63. (**A-D**) Confocal fluorescence micrographs of (A, B) 293F cells and (C, D) 293F/AP2M1^-/-^ cells that had been chilled, incubated with fluorescent antibodies specific for (green) CD63 and (pink) CD9, washed, warmed to 37°C, and then fixed after (A, C) 0 minutes at 37°C or (B, D) 30 minutes at 37°C. Bar, 10 um. (**E**) Histograms of flow cytometry data collected from 293F and 293F/AP2M1^-/-^ cells that had been chilled and surface stained with fluorescently tagged antibodies specific for CD63, CD9, and CD81. (**F**) Immunoblot of cell and exosome fractions collected from 293F and AP2M1^-/-^ cells, probed with antibodies specific for CD63, CD81, CD9, Hsp90, and AP2M1. (**G**) Bar graph showing the mean relative exosomal secretion of CD63 in 293F and 293F/AP2M1^-/-^ cells, +/- s.e.m., calculated from immunoblot data. Student’s t-test revealed a *p* value <0.005. (**H**) Immunoblot of exosome fractions collected from 293F cells cultured in medium lacking or containing latrunculin A, probed with antibodies specific for CD63, CD9, and CD81. (**I**) Bar graph showing the mean CD63/CD81 ratio +/- s.e.m. in exosomes of 293F cells incubated in the absence or presence of latrunculin A. Student’s t-test revealed a *p* value of 0.0028. All experiments were performed a minimum of three times. Dat is from three independent trials.

Previous studies have established that actin polymerization plays a critical role in protein endocytosis by providing a driving force for membrane invagination, that latrunculin A inhibits actin polymerization by sequestering actin monomers, and that hat latrinculin A also inhibits protein endocytosis(*40-44*). In light of these considerations, latrunculin A should also induce a selective increase in the exosomal secretion of CD63. To test this prediction, we grew 293F cells in media lacking or containing latrunculin A, collected their exosomes, and interrogated them by immunoblot for CD63, CD81, and CD9. This revealed that latrunculin increased the exosomal secretion of CD63 by ∼7-fold but had no effect on the exosomal secretion of CD81 and only a minor effect on the exosomal secretion of CD9 (***Fig. 3H, I***), providing additional evidence that inhibiting endocytosis is sufficient to selectively induce the exosomal secretion of CD63.

### Endocytosis signals inhibit the exosomal secretion of cargo proteins in *cis* and in *trans*

The antagonism between endocytosis and exosome biogenesis is consistent with our previous discoveries that endosome membrane anchors are unable to induce the exosomal secretion of cargo proteins(*33*), that appending an endocytosis signal to the C-terminus of CD9 greatly reduced its exosomal secretion from the cell(*3, 4*), and that elimination of CD63’s endocytosis signal greatly increased its exosomal secretion from the cell(*3, 4*) (a result later confirmed by others(*11*)). Here we extend these studies by exploring the post-endocytosis fates of exosome cargoes, the effect of cargo protein expression level, or the fact that endocytosis signal-containing proteins can saturate AP-2 and inhibit protein endocytosis(*45, 46*). Specifically, we used Cas9-mediated gene editing to generate a CD9^-/-/-^ 293F cell line (***fig. S6***), converted it to a Tet-on cell line (FtetZ/CD9^-/-/-^), and then created derivatives of this FtetZ/CD9^-/-/-^ cell line that carry doxycycline-regulated transgenes designed to express WT CD9 or mutant forms of CD9, including CD9-YQRF, which does not bind syntenin but has high affinity for the mu2 subunit of AP-2(*35*), CD9-YQTI, which also does not bind syntenin but binds the mu2 protein and also the mu3 subunit of AP-3(*35*), CD9-YEVM, which binds syntenin and also AP-2 and AP-3(*24, 35*), and CD9-AEMV, which lacks the motifs for binding syntenin, AP-2, or AP-3.

Although qRT-PCR experiments showed that level of transgene-encoded CD9 mRNA was similar in all cell lines (***fig. S7***), immunoblot experiments showed that the amount of CD9 protein was not (***Fig. 4A***). Specifically, CD9 and CD9-AEMV were readily detected in both cell and exosome lysates whereas we were unable to detect the CD9-YQRF, CD9-YQTI, or CD9-YEVM in either cells or exosomes. Furthermore, addition of the vacuolar ATPase (V-ATPase) inhibitor bafilomycin A partly restored the abundance of these proteins (***fig. S8***), indicating that CD9-YQRF, CD9-YQTI, and CD9-YEVM were being degraded by the plasma membrane-to-lysosome trafficking pathway(*47-49*). These results demonstrate that endocytosis inhibits the exosomal secretion of cargo proteins by targeting them for destruction, as well as by removing them from the primary site of exosome biogenesis at the plasma membrane.

**Figure 4.**
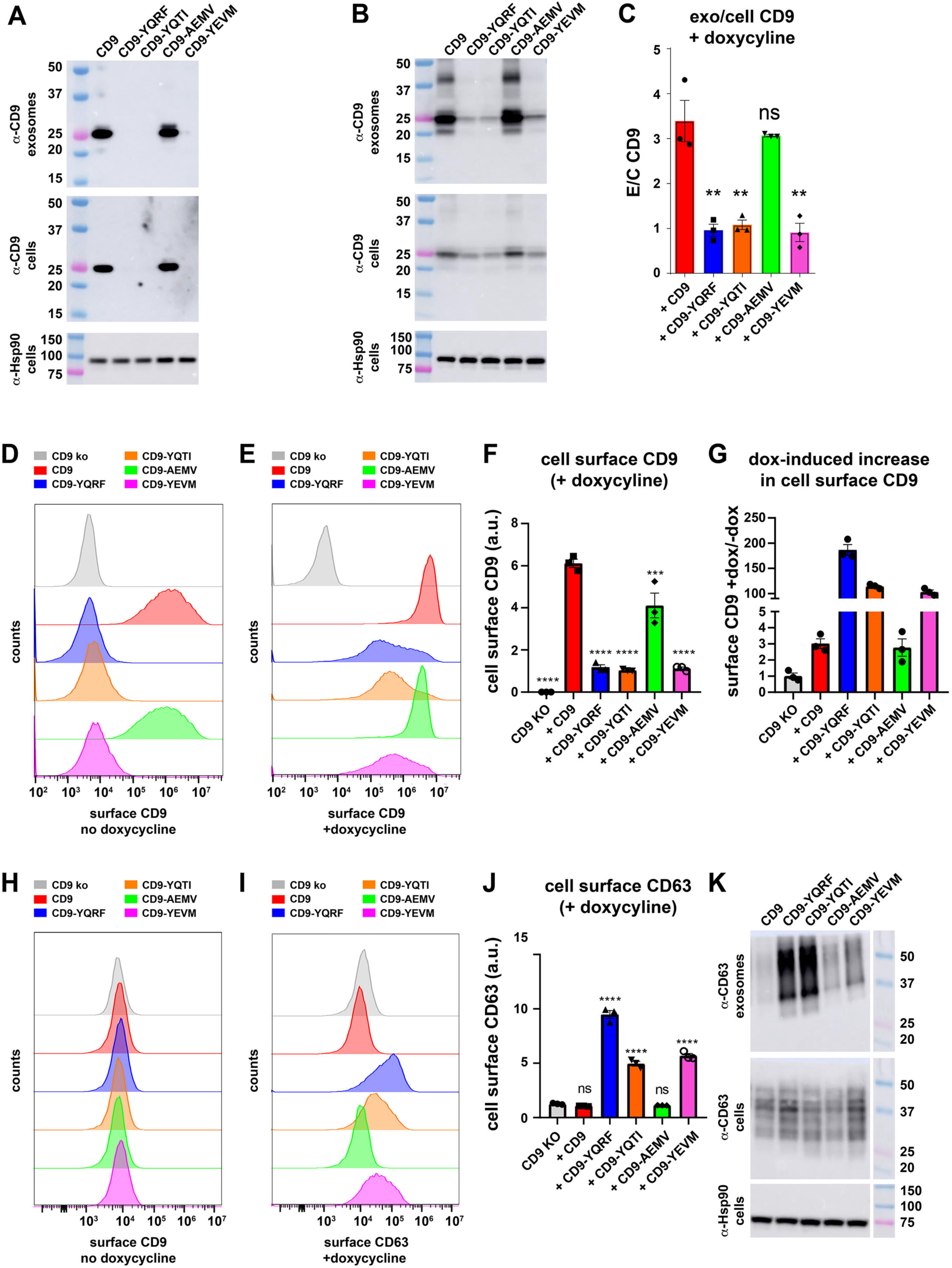
Endocytosis of CD9 inhibits its exosomal secretion and induces its lysosomal destruction. (**A, B**) Immunoblots of (upper panel) exosome lysates and (lower panels) cell lysates from (A) uninduced and (B) doxycycline-induced FtetZ/CD9^-/-/-^ cells carrying TRE3G-regulated transgenes that express WT CD9, CD9-YQRF, CD9-YQTI, CD9-AEMV, or CD9-YEVM, probed with antibodies to CD9, with cell lysates also probed with antibodies to Hsp90. MW markers are in kDa. (**C**) Bar graphs showing the relative exosomal secretion of each CD9 protein from doxycycline-induced cell lines (amount of protein in exoxomes/amount of protein in cell). ** denotes an ANOVA p value of <0.005. (**D, E**) Histograms of anti-CD9 flow cytometry of (D) uninduced or (E) doxycycline-induced FtetZ/CD9^-/-/-^ cells and FtetZ/CD9^-/-/-^ cells carrying TRE3G-regulated transgenes that express WT CD9, CD9-YQRF, CD9-YQTI, CD9-AEMV, or CD9-YEVM. (**F**) Bar graph showing the mean +/- s.e.m. of cell surface CD9 staining, with **** and *** denoting ANOVA *p* values of <0.0005 and <0.005, respectively. (**G**) Bar graph showing the mean +/- s.e.m. of the fold difference in cell surface staining for CD9 in doxycycline-induced/uninduced cells. Data is from three independent trials. (**H**, **I**) Histograms of anti-CD63 flow cytometry of (H) uninduced or (I) doxycycline-induced FtetZ/CD9^-/-/-^ cells and FtetZ/CD9^-/-/-^ cells carrying TRE3G-regulated transgenes that express WT CD9, CD9-YQRF, CD9-YQTI, CD9-AEMV, or CD9-YEVM. (**J**) Bar graph showing the mean +/- s.e.m. of cell surface CD63 staining, with **** denoting ANOVA *p* values of <0.0005. (**K**) Immunoblots of (upper panel) exosome lysates and (lower panels) cell lysates from doxycycline-induced FtetZ/CD9^-/-/-^ cells carrying TRE3G-regulated transgenes that express WT CD9, CD9-YQRF, CD9-YQTI, CD9-AEMV, or CD9-YEVM, probed with antibodies to CD63, with cell lysates also probed with antibodies to Hsp90. MW markers are in kDa.

To determine whether the level of CD9 expression has any effect on the dynamics of CD9 stability or exosomal secretion, we repeated these experiments on cell and exosome lysates collected from doxycycline-induced cell cultures. As expected, CD9 and CD9-AEMV were even more abundant in both cell and exosome lysates (***Fig. 4B***). Furthermore, we found that high-level expression of CD9-YQRF, CD9-YQTI, or CD9-YEVM led to their accumulation in the cell to the point where they could be easily detected, and also led to their exosomal secretion from the cell. This allowed us to calculate the relative exosomal secretion of all five CD9 proteins, which proved to be ∼3-fold higher for CD9 and CD9-AEMV than for CD9-YQRF, CD9-YQTI, and CD9-YEVM (n = 3, *p* <0.005), demonstrating once again that endocytosis inhibits the vesicular secretion of exosome cargo proteins (***Fig. 4C***).

These data also raised the question of why the plasma membrane-to-lysosome trafficking/destruction of CD9-YxxΦ proteins was inhibited by their doxycycline-induced expression. This pathway has multiple steps that are potentially subject to saturating inhibition (clathrin-mediated endocytosis, uncoating of endocytic vesicles, fusion of these vesicles with endosomes, loading of endocytosed CD9 into ILVs, delivery of CD9-ILVs to lysosomes, and lysosomal hydrolysis of CD9-ILV lipids and proteins). However, the most notable of these is endocytosis, as previous studies have already shown that AP-2-mediated endocytosis is inhibited by high-level expression of endocytosis signal-containing proteins(*45, 46*). To explore this possibility, we used flow cytometry to measure the cell surface expression of each CD9 protein at both their baseline and induced levels of expression.

In the absence of doxycycline, CD9 and CD9-AEMV were abundant at the plasma membrane, whereas anti-CD9 staining for CD9-YQRF, CD9-YQTI, and CD9-YEVM was minimal, only slightly higher than the background staining observed for F/CD9^-/-/-^ control cells (***Fig. 4D***). In presence of doxycycline, the plasma membrane levels of CD9 and CD9-AEMV rose by ∼3-fold, consistent with the higher level of exosomal secretion seen for the doxycycline-induced cultures of these cell lines. However, the biggest impact of doxycycline-induced expression was the ∼100-fold increase in the cell surface abundance of CD9-YQRF, CD9-YQTI, and CD9-YEVM (***Fig. 4D-G***). The end result of this ∼100-fold increase was that their plasma membrane abundance was only ∼5-fold less than WT CD9 and CD9-AEMV (n = 3, *p* < 0.0005), raising the possibility that the expression-induced exosomal secretion of these CD9-YxxΦ proteins (***Fig. 4B***) was caused by their plasma membrane accumulation and direct budding from the cell surface.

If high-level expression of CD9-YQRF, CD9-YQTI, and CD9-YEVM was inhibiting AP-2-mediated endocytosis, this inhibition should be reflected in an expression-dependent, CD9-YxxΦ- specific increase in the plasma membrane abundance of CD63. At baseline levels of CD9 transgene expression, the cell surface abundance of CD63 was the same in F/CD9^-/-/-^ control cells as in each of the CD9-expressing cell lines (***Fig. 4H***). However, the doxycycline-induced expression of CD9-YQRF, CD9-YQTI, or CD9-YEVM led to a pronounced increase in the cell surface abundance of CD63, 10-fold in the case of CD9-YQRF and 5-fold in the case of CD9-YQTI or CD9-YEVM, while high-level expression of WT CD9 or CD9-AEMV had no effect on the cell surface abundance of CD63 (***Fig. 4H-J***). Given that similar increases in plasma membrane accumulation of CD63 are associated with increased exosomal secretion of CD63, we interrogated cell and exosome fractions from doxycycline-induced cultures by immunoblot for CD63, observing that high level expression of CD9-YQRF or CD9-YQTI, and to a lesser extent CD9-YEVM, induced the exosomal secretion of CD63 (***Fig. 4K***).

### CD63 is an expression-dependent regulator of AP-2-mediated endocytosis and exosome content

The preceding results demonstrate a complex interplay between the expression level of YxxΦ- containing exosome cargo proteins, their endocytosis, their exosomal secretion, and the endocytosis and exosomal secretion of other YxxΦ-containing proteins. To determine whether CD63 displayed a similar complexity of effects on the sorting and exosomal secretion of YxxΦ-containing exosome cargo proteins, we first tested whether it display an expression-dependent inhibition of its own endocytosis. Towards this end, we created FtetZ/CD63^-/-^ cells, a Tet-on derivative of the previously described 293F CD63^-/-^ cell line(*50*), and then generated a derivative of FtetZ/CD63^-/-^ cells that carries a doxycycline-regulated, *TRE3G*-CD63 transgene, the FtetZ/CD63^-/-^::*TRE3G*-CD63 cell line. These cells were seeded onto glass coverslips, incubated overnight in normal medium or medium supplemented with doxycycline, chilled to 4°C, incubated with fluorescently-labeled antibodies specific for CD63, washed, and then fixed either immediately (t = 0) or after a 30 minutes-long incubation at 37°C (t = 30). When these cells were examined by confocal fluorescence microscopy, we observed that CD63 was endocytosed by the uninduced cells but not by the doxycycline-induced cells (***Fig. 5***), demonstrating that high-level expression of WT CD63 inhibits its own endocytosis.

**Figure 5.**
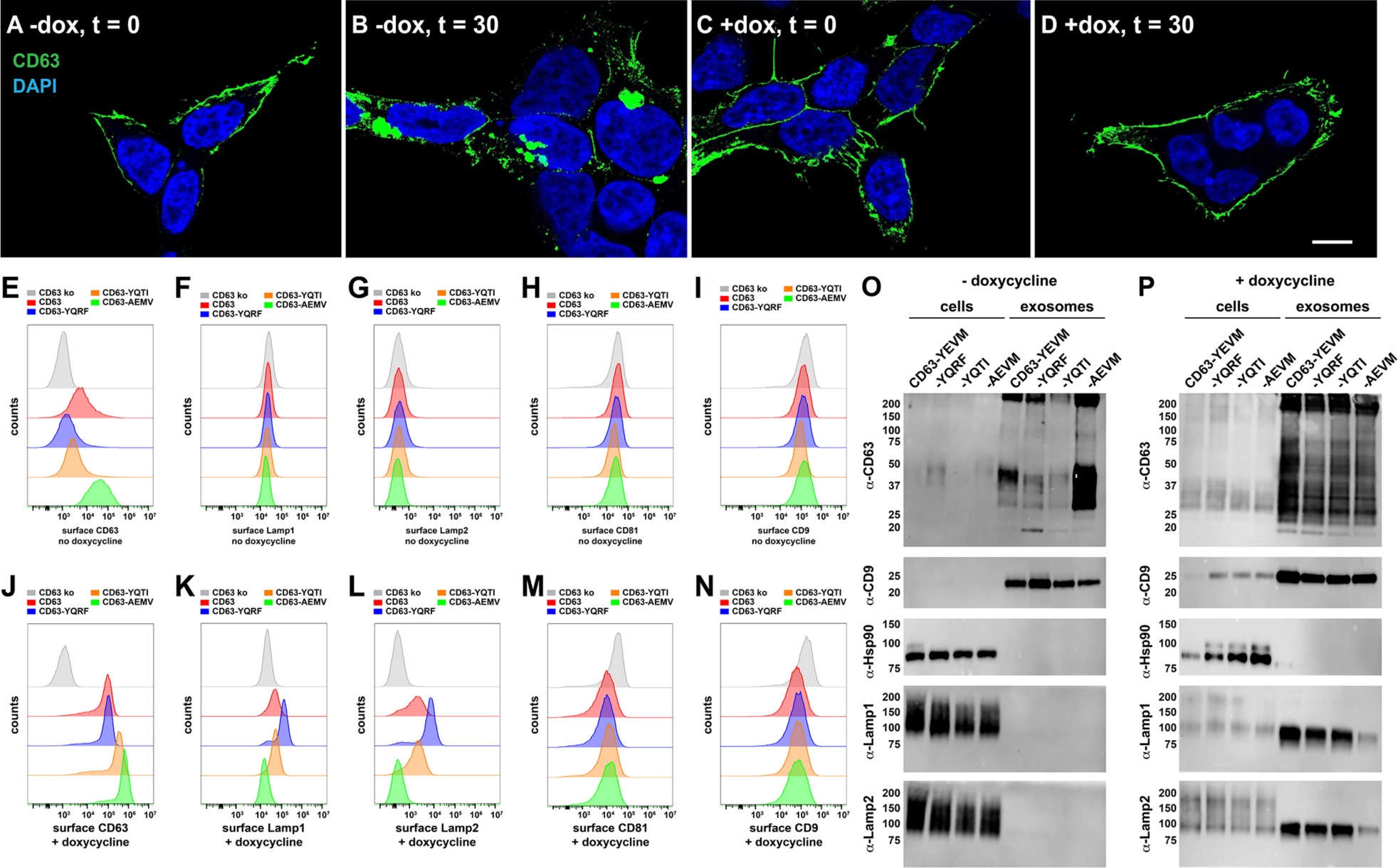
Endocytosis inhibits the exosomal secretion of CD63 while CD63 inhibits endocytosis. (**A-D**) Confocal fluorescence micrographs of FtetZ/CD63^-/-^ ::CD63 cells grown in the (A, B) absence or (C, D) presence of doxycycline that had been chilled, incubated with a fluorescent antibody specific for (green) CD63, washed, fixed after (A, C) 0 minutes at 37°C or (B, D) 30 minutes at 37°C, then stained with DAPI. Bar, 10 um. (**E-I**) Histograms of flow cytometry data collected on uninduced (grey) FtetZ/CD63^-/-^ cells and FtetZ/CD63^-/-^ cells carrying TRE3G-regulated transgenes that express (red) WT CD63, (blue) CD63-YQRF, (orange) CD9-YQTI, or (green) CD63-AEMV, stained for cell surface (E) CD63, (F), Lamp1, (G) Lamp2, (H) CD81, or (I) CD9. (**J-N**) Histograms of flow cytometry data collected on doxycycline-induced (grey) FtetZ/CD63^-/-^ cells and FtetZ/CD63^-/-^ cells carrying TRE3G-regulated transgenes that express (red) WT CD63, (blue) CD63-YQRF, (orange) CD9-YQTI, or (green) CD63-AEMV, stained for cell surface (J) CD63, (K), Lamp1, (L) Lamp2, (M) CD81, or (N) CD9. (**O, P**) Immunoblot of cell and exosome lysates collected from (O) uninduced and (P) doxycycline-induced FtetZ/CD63^-/-^ cells carrying TRE3G-regulated transgenes that express WT CD63, CD63-YQRF, CD9-YQTI, or CD63-AEMV. Blots were probed with antibodies specific for CD63, CD81, CD9, Lamp1, and Lamp2. MW size markers are listed in kDa. Data is from three independent trials.

We next tested the effect of CD63 endocytosis signal substitutions and expression level on the cell surface abundance of CD63 and other proteins. Towards this end, we created three more FtetZ/CD63^-/-^::*TRE3G*-CD63-like cell lines designed to express CD63-YQRF (binds AP-2 but not syntenin), CD63-YQTI (binds AP-2 and AP-3 but not syntenin) or CD63-AEMV, which lacks an endocytosis signal. These cells, together with FtetZ/CD63^-/-^ cells and FtetZ/CD63^-/-^::*TRE3G*-CD63 cells, were then grown in normal medium, chilled, stained with antibodies, and examined by flow cytometry. In the uninduced state, the cell surface abundance of CD63 proteins was determined by the sequence of their C-terminal endocytosis signal, with the non-endocytosed CD63-AEMV protein accumulating at the plasma membrane at levels that were 10-fold higher than that of WT CD63 and 20-fold higher than that of CD63-YQRF or CD63-YQTI (***Fig. 5E***). These results indicate that this baseline level of CD63-YxxΦ protein expression was too low to saturate AP-2 and inhibit their endocytosis. In support of this interpretation, the cell surface levels of Lamp1, Lamp2, CD81, and CD9 were the same in FtetZ/CD63^-/-^ cells and in each of the four CD63-expressing cell lines (***Fig. 5F-I***).

In parallel with these studies, we also performed flow cytometry on doxycycline-induced cell cultures as a way assess the impact of increasing the expression of CD63. These experiments revealed that high level expression led to the plasma membrane accumulation of all four forms of CD63, with the cell surface abundance of CD63-AEMV increasing by ∼10-fold while the cell surface of CD63-YxxΦ proteins increased even more, ∼20-fold for WT CD63 and ∼50-fold for CD63-YQRF and CD63-YQTI (***Fig. 5J***). These results indicate that high level expression of CD63-YxxΦ proteins had saturated the AP-2 machinery. Consistent this interpretation, we found that cell surface abundance of Lamp1 and Lamp2 were both increased in cells expressing high levels of WT CD63, CD63-YQRF or CD63-YQTI (***Fig. K, L***), with cell surface Lamp1 increasing 2-fold in cells expressing WT CD63 or CD63-YQTI and 5-fold in cells expressing CD63-YQRF, while cell surface Lamp2 increased 20-fold in cells expressing WT CD63 or CD63-YQTI and 50-fold in cells expressing CD63-YQRF. Flow cytometry also revealed that doxycycline-induced expression of all four CD63 proteins reduced the plasma membrane abundance of both CD81 and CD9 by ∼2-fold to 4-fold, an effect that was unrelated to the presence of an AP-2-binding YxxΦ motif (***Fig. 5M, N***), which might reflect a competition in the endoplasmic reticulum-to-plasma membrane trafficking of exosomal tetraspanins.

To determine whether the plasma membrane accumulation of CD63, Lamp1, and Lamp2 bore any relation to their exosomal secretion from the cell, we collected cell and exosome fractions from uninduced and induced cell cultures, then interrogated them by immunoblot. In uninduced cells, the differences in plasma membrane abundance between WT CD63, CD63-YQRF, CD63-YQTI, and CD63-AEMV accurately predicted their relative loading into exosomes, as the form of CD63 that lacks an endocytosis signal (CD63-AEMV) displayed the highest exosomal secretion, CD63-YQRF and CD63-YQTI displayed the lowest degree of exosomal secretion, and WT CD63 displayed an intermediate level of exosomal secretion, presumably because its endocytosis signal is masked by syntenin (***Fig. 5O***). These cells displayed little if any exosomal secretion of either Lamp1 or Lamp2, consistent with the flow cytometry data showing that their cell surface expression was unaffected by the baseline level of CD63 protein expression. In contrast, doxycycline-induced expression led to the efficient exosomal secretion of all four CD63 proteins, regardless of whether they did or did not carry a functional endocytosis signal (***Fig. 5P***). Furthermore, we found that high-level expression of WT CD63, CD63-YQRF, or CD63-YQTI induced a dramatic increase in the exosomal secretion of Lamp1 and Lamp2, providing yet more evidence that high-level expression of CD63 inhibits AP-2-mediated endocytosis, and that plasma membrane accumulation is a precursor to exosomal secretion.

### Co-regulation of syntenin, CD63, and the plasma membrane biogenesis of CD63 exosomes

CD63 expression varies from >3,000 normalized transcripts per million (nTPM) in airway epithelial cells to less than 10 nTPMs in neurons (***Table S1***)(*51*), marking CD63 as one of the most highly expressed genes and also one of the most variably expressed human genes. This raises the question of how cells manage CD63’s expression-dependent inhibition of AP-2-mediated endocytosis. One potential mechanism is that cells coordinately express both CD63 and syntenin, a scenario that would allow syntenin to suppress CD63’s inhibition of AP-2. In support of this hypothesis, we found that CD63 and syntenin displayed a strong positive correlation in expression across 44 distinct sets of normal and cancer tissues (***Fig. S9***). This coordinate expression was also evident at the protein level, as syntenin expression was highest in SK-MEL-5 cells (2373 nTPM), moderate in 293 cells (249 nTPM), and lowest in Daudi cells (87 nTPM) (***Table S1***, ***Fig. 6A***).

**Figure 6.**
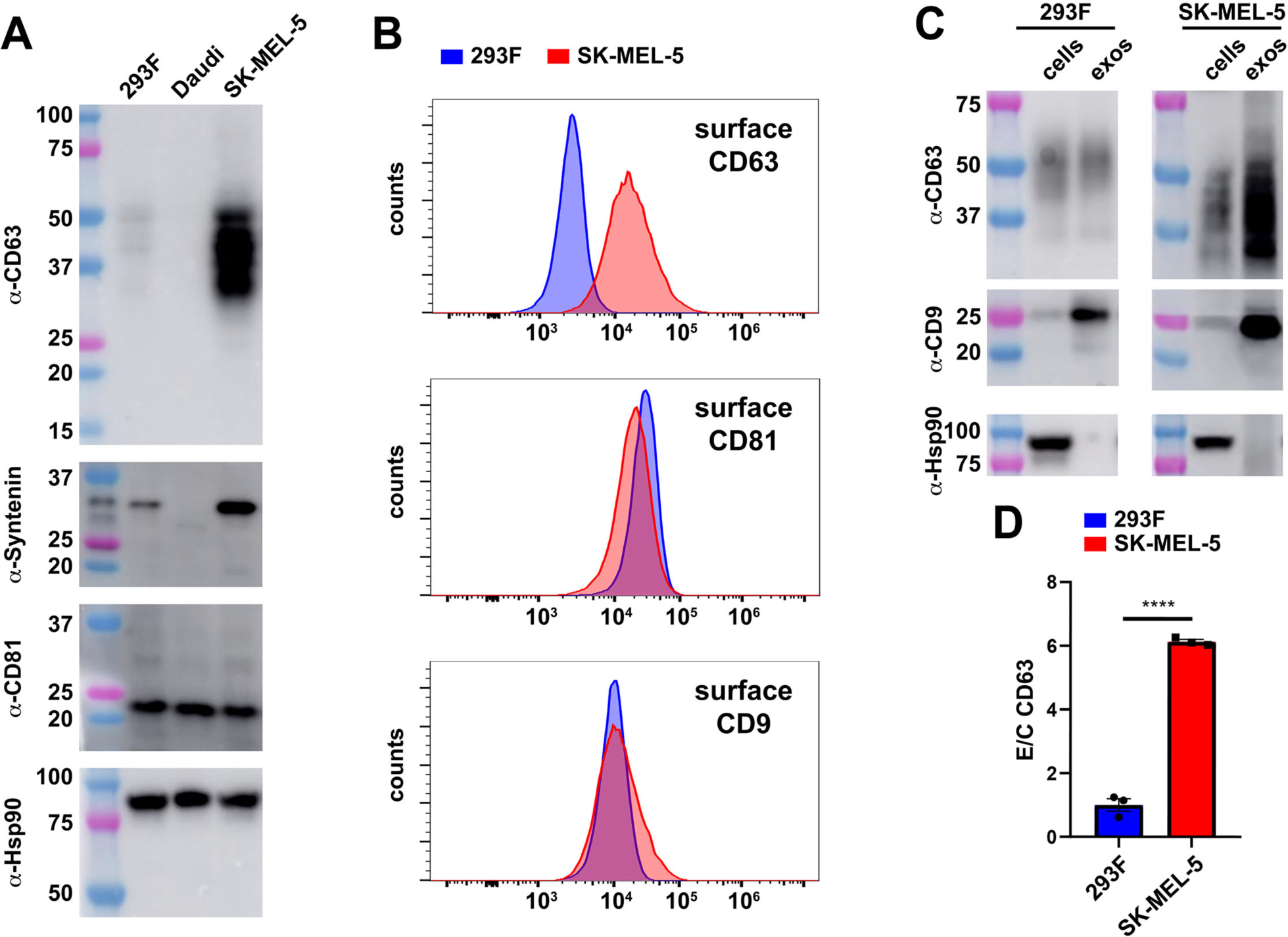
Natural elevation of CD63 expression also impairs endocytosis and induces its exosomal secretion. (**A**) Immunoblot of cell lysates collected from 293F, Daudi, and SK-MEL-5 cells using antibodies specific for CD63, syntenin, CD81, and Hsp90. MW size markers are listed in kDa. (**B**) Histograms of cell surface flow cytometry of 293F and SK-MEL-5 cells, stained with antibodies specific for CD63, CD81, and CD9. (**C**) Immunoblot of cell and exosomes lysates collected from 293F and SK-MEL-5 cells, probed with antibodies to CD63, CD9, and Hsp90. Bar graph showing the efficiency of CD63 budding (exosome/cell ratio) from 293F and SK-MEL-5 cells, with bar height representing the mean and error bars the s.e.m. (error bars). Data is from three independent trials.

Given that high-level expression of either syntenin or CD63 inhibit CD63’s endocytosis and induce its plasma membrane accumulation and exosomal secretion, the high-level expression of both CD63 and syntenin in SK-MEL-5 cells predicts that this cell line should display significantly higher cell surface accumulation and exosomal secretion of CD63 than 293F cells. Flow cytometry confirmed the first half of this prediction, as SK-MEL-5 cells accumulated CD63 to nearly 10-times the level seen in 293F cells, even though both cell lines had similar levels of cell surface CD81 and CD9 (***Fig. 6B***). Furthermore, when we collected cell and exosome fractions from SK-MEL-5 and 293F cells and interrogated them by immunoblot, we found that SK-MEL-5 cells loaded CD63 into exosomes at ∼6-fold higher efficiency than 293F cells (***Fig. 6C, D***), nearly the same increase in exosomal secretion that we observed for CD63 proteins lacking their endocytosis signal(*3, 4*).

## Discussion

The results presented here show that syntenin’s primary role in exosome biogenesis is to inhibit CD63 endocytosis, drive the plasma membrane accumulation of CD63, and thereby induce its loading into plasma membrane-derived exosomes. In support of these conclusions, we also showed that endocytosis is generally antagonistic to the exosomal secretion of exosome cargo proteins, that inhibitors of endocytosis are sufficient to induce the plasma membrane accumulation and exosomal secretion of CD63, and that high-level expression of CD63 inhibits AP-2-mediated endocytosis and induces the plasma membrane accumulation and exosomal secretion of CD63 and other lysosome membrane proteins, including Lamp1 and Lamp2. Furthermore, we showed that the indirect mechanism of syntenin-induced CD63 budding was independent of Alix. Taken together, these data support and extend the shared, stochastic hypothesis of exosome biogenesis(*3, 4, 6*) in which exosome cargo proteins bud along the continuum of plasma and endosome membranes, though primarily from the plasma membrane (***Fig. 7***).

**Figure 7.**
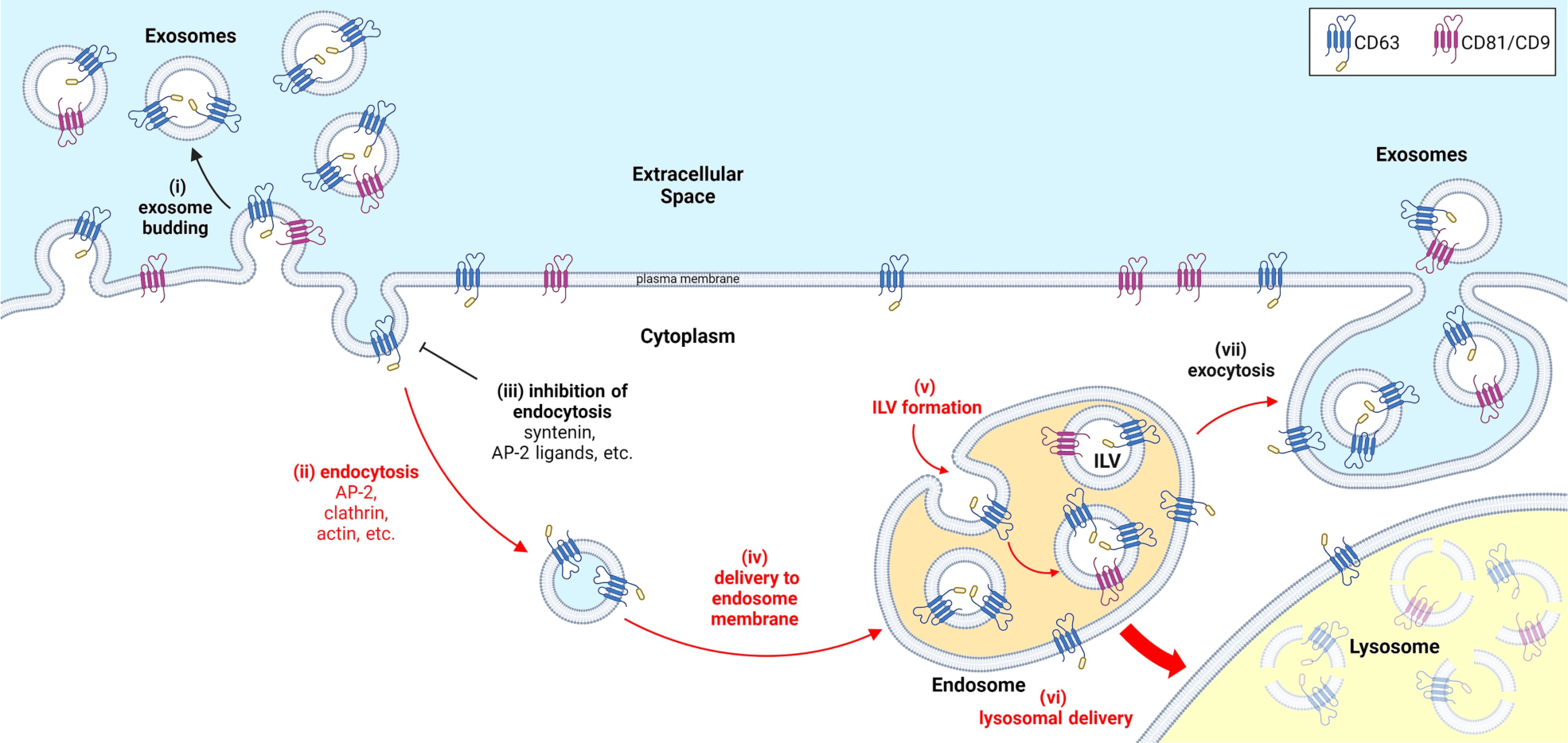
The shared, stochastic model of exosome biogenesis, updated. The plasma membrane is (***i***) the primary site of exosome biogenesis. Thus, the localization of exosome cargoes to the plasma membrane is a primary determinant of their exosomal secretion, which is why (***ii***) endocytosis is a potent inhibitor of a protein’s exosomal secretion. Under this model, any factor that (***iii***) inhibits the endocytosis of an exosome cargo will induce its exosomal secretion from the cell, which is why so many inhibitors of CD63 endocytosis induce its exosomal secretion from the cell (mutation of AP2M1, inhibition of actin polymerization, saturating inhibition of AP-2, etc.). This model also helps us understand the fate of endocytosed exosome cargo proteins, most of which are which are (***iv***) delivered to endosome membranes where they are (***v***) loaded into nascent ILVs. Many of these ILVs are (***vi***) delivered to lysosomes and destroyed by lysosomal hydrolases, completing a plasma membrane-to-lysosome degradative pathway. However, defects in endosome-lysosome fusion, blockade of lysosome hydrolases, repair of plasma membrane tears, or other inducers of endolysosomal exocytosis can (***vii***) result in ILV secretion. Although this endocytosis/exocytosis route of exosome biogenesis is commonly slow, saturable, and secondary to the plasma membrane route of exosome biogenesis, it can mediate an *en bloc* release of whatever stockpiles of ILVs happen to reside in the cell at any given time, and can lead to significant exosome release in response to specific signals.

### Exosome cargo proteins bud primarily from the plasma membrane

Exosome cargo proteins bud best when they are localized to the plasma membrane (***Fig. 7i***). This was facet of exosome biogenesis was first demonstrated by us in 2011 in the T-cell leukemia cell line Jurkat (*31, 33*), and more conclusively by us again in 2019(*4*) and 2022(*3*) in the human non-cancerous cell line HEK293, the mouse fibroblast cell line NIH3T3, and in human primary fibroblasts, and then again in this report in 293F cells, NIH3T3 cells, Hela cells, and SK-MEL-5 cells. Furthermore, the empirical support for this conclusion includes a wide array of experimental observations, including the fact that plasma membrane-localized exosome cargoes like CD81 and CD9 are secreted from the cell in exosomes at far greater efficiency than endocytosed cargoes like CD63, that inhibition of CD63’s endocytosis by syntenin is sufficient to induce CD63’s plasma membrane accumulation and exosomal secretion, that other inhibitors of endocytosis are also sufficient to induce the selective plasma membrane accumulation and exosomal secretion of CD63, that appending endocytosis signals to CD9 dramatically reduce its exosomal secretion, that removing the endocytosis signal from CD63 dramatically induced its exosomal secretion, and that inhibiting AP-2-mediated endocytosis also induced the plasma membrane accumulation and exosomal secretion of Lamp1 and Lamp2.

### Endocytosis inhibits the exosomal secretion of exosome cargo proteins

In addition to its direct empirical support, the primary role of the plasma membrane in exosome biogenesis is consistent with the fact that the plasma membrane has an immense surface area (∼2000 μm^2^), that every plasma membrane vesicle budding event generates an exosome, and that (***Fig. 7ii***) endocytosis inhibits the exosomal secretion of exosome cargo proteins. The evidence for the antagonistic effect of endocytosis on exosome biogenesis is extensive, and includes the reduced exosomal secretion in cells knocked out for syntenin, the reduced exosomal secretion of CD9-YxxF proteins, and the reduced exosomal secretion of CD63-YxxΦ proteins. Furthermore, this paper also established that (***Fig. 7iii***) inhibitors of endocytosis induce the plasma membrane accumulation and exosomal secretion of CD63, a result we observed for cells expressing high levels of syntenin, mutated in AP2M1, incubated with latrunculin, or expressing AP-2-inhibiting levels of CD9-YxxF proteins or CD63. Furthermore, these relationships between endocytosis, inhibition of endocytosis, plasma membrane accumulation, and exosomal secretion were also evident in the behavior of Lamp1 and Lamp2 in cells that expressed AP-2-inhibiting levels of CD63.

### Fates of endocytosed exosome cargo proteins

Under our working hypothesis of exosome biogenesis (***Fig. 7***), endocytosis of an exosome cargo protein leads to (***Fig. 7iv***) its delivery to the limiting membrane of endosomes, (***Fig. 7v***) its loading into nascent intraluminal vesicles (ILVs), and in most cases, (***Fig. 7vi***) its ILV-mediated delivery to lysosomes, and subsequent destruction. This canonical, plasma membrane-to-lysosome trafficking pathway appears to explain the fate of endocytosis signal-containing forms of CD9, as these proteins were only detectable when the pathway was inhibited, either by addition of the V-ATPase inhibitor bafilomycin or by saturating inhibition of AP-2 induced by the high-level expression of these CD9-YxxΦ proteins. In contrast to CD9, endocytosis of CD63 was not correlated with its efficient turnover, and thus, the plasma membrane-to-lysosome trafficking of different exosome cargo proteins appears to proceed at distinct efficiencies. As for the mechanistic basis of these differences, we do not currently know. However, some possibilities are that CD63 may be delivered to endosomes that are more refractory to fusion with lysosomes, and/or that CD63, as one of the most abundant lysosome membrane proteins known, is particularly resistant to lysosomal proteases.

Regardless of these differences, there is no doubt that the plasma membrane-to-lysosome trafficking pathway is multi-step, slow, and saturable. This results in the presence of a steady-state stockpile of yet-to-be-degraded ILVs, some or all of which can be (***Fig. 7vii***) released as exosomes whenever an ILV-containing endosome or lysosome fuses with the plasma membrane. In the present paper, our experiments were designed to detect the exosomal secretion of cargo proteins by this endocytosis/exocytosis route of exosome biogenesis, but in all cases our data indicated that its contribution to exosome biogenesis was minimal (***Table S2***). This endocytosis/exocytosis pathway of exosome biogenesis can, however, be induced by any of several triggers of endolysosomal exocytosis, including activators of cytoplasmic calcium (plasma membrane tears, ionomycin, etc.(*52*)), lysosome poisons (e.g. bafilomycin A(*53*)), virus-encoded de-acidifying proteins(*54*), and mutational inactivation of genes required for endosome-lysosome fusion (e.g. VPS39(*55*)). Less clear is whether the increased exosomal secretion induced by these triggers is coming primarily from ILV secretion or by increased exosome budding directly from the plasma membrane.

### Relevance to exosome engineering

In addition to being the simplest explanation of the available data (***Table S2*** and (*3*)), the shared, stochastic model of exosome biogenesis (***Fig. 7***), and particularly its foundational observation that the most highly-enriched exosome cargoes all bud best when localized to the plasma membrane(*3, 4, 6, 11, 31, 33*), draw a clear and simple roadmap for exosome engineering. In brief, this roadmap shows that nearly any protein can be loaded into exosomes by maximizing just three variables: its expression, its delivery to the plasma membrane, and its loading into nascent, plasma membrane budding vesicles. As for whether it’s possible to engineer highly expressed proteins into and through the endocytosis/exocytosis route of exosome biogenesis, our data indicate that it’s unlikely, in part because endocytosed cargoes are destroyed by the plasma membrane-to-lysosome trafficking pathway, and in part because high-level expression of an endocytosis signal-containing cargo will block its endocytosis and induce its direct budding from the plasma membrane. In short, any attempt to engineer the endocytosis/exocytosis pathway will undermine its own goals, induce cargo budding from the plasma membrane, and likely disrupt exosome content further by inducing the plasma membrane accumulation of otherwise endocytosed proteins.

### Implications for CD63-based models of exosome biology

In addition to their implications for exosome biogenesis, our data reveal a previously unrecognized flaw in studies that make use of CD63 overexpression(*11, 56-66*). Specifically, we demonstrated that high-level expression of CD63 inhibits its own endocytosis, inhibits the endocytosis of other proteins, drives its direct budding from the plasma membrane, and drives the aberrant exosomal secretion of Lamp1, Lamp2, and likely other AP-2-binding proteins as well. Not surprisingly, evidence for this can be seen in the data of some earlier papers. For example, the CD63-induced inhibition of AP-2-mediated endocytosis and resulting exosomal secretion of CD63, Lamp1, Lamp2, and CD9 is also evident in the data of Mathieu et al.(*11*), which reported that high-level expression of CD63 led to its co-budding from the cell with the plasma membrane marker CD9, and also induced the exosomal secretion of Lamp1 and Lamp2. Furthermore, the previously unexplained lethality of CD63-GFP transgenic mice(*56*) takes on greater significance, as it adds independent genetic evidence that CD63 overexpression systems have unintended and adverse effects on cells and animals. In light of these considerations, we urge caution when using CD63-based experimental models(*11, 56-66*), and recommend instead the use of models based on the more highly-enriched and non-endocytosed cargoes CD81 and CD9.

### Mechanism and logic in exosome nomenclature

In contrast to the shared, stochastic model of exosome biogenesis (***Fig. 7***), others have proposed that CD63 exosomes are generated exclusively by endocytosis/exocytosis route of exosome biogenesis (*11-23*). Proponents of this ‘endosome-dependent’ hypothesis also assert that it’s possible to tell the difference between exosome-sized vesicles that are generated by direct budding from the plasma membrane versus those that arise by endocytosis/exocytosis, and moreover, that these two classes of vesicles should be referred to with different names The data presented here refute all three of these assertions. In experiment after experiment, our data support the shared, stochastic hypothesis of exosome biogenesis (***Fig. 7***) and argue strongly against the idea that exosomes are made solely or even primarily by the endocytosis/exocytosis pathway (***Table S2***). Furthermore, the data presented here and in previous reports(*3, 4*) shows definitively that exosome cargo proteins bud from plasma and endosome membranes in exosomes of the same size, topology and overlapping set of proteins. Thus, there is no way to determine where an exosome arose once it has left the cell, and therefore no rational basis for a dichotomous nomenclature that pretends what cannot be done. We therefore urge the EV field to adopt a definition of exosomes that can be experimentally defined, such as *small secreted vesicles of ∼30-150 nm that have the same topology as the cell and are enriched in exosome cargo proteins*(*6, 67, 68*). This simple definition accurately reflects the underlying biology of exosome cargo protein trafficking and exosome biogenesis, respects the fact that the origin membrane of individual exosomes cannot be determined, and is consistent with the complex interactions between exosome protein budding, exosome biogenesis, endocytosis, lysosomal protein trafficking and endolysosomal exocytosis (***Fig. 7***).

## Materials and Methods

### Plasmids

The plasmid pJM1463 is based on a pS series plasmid(*69*) and carries a bicistronic ORF encoding rtTAv16-2a-BleoR downstream of the spleen focus forming virus (SFFV) transcriptional control region. The plasmids pYA128, YA129, and pYA130 are Sleeping Beauty transposons based on pITRSB(*69*) that carry two genes: (***i***) an EFS-PuroR gene and (***ii***) a TRE3G regulated gene designed to express syntenin, N100syntenin, or syntenin C23, respectively. The plasmids pCG606, pCG732, pCG733, pCG734, and pCG607 are Sleeping Beauty transposons based on pITRSB that carry two genes: (***i***) an EFS-HygR gene and (**ii**) a TRE3G regulated gene designed to express codon optimized CD9 ORFs that encode WT CD9, CD9-YQRF, CD9-YQTI, CD9-AEMV, and CD9-YEVM, respectively. The plasmids pCG602, pMG9, pCG604, and pMG10 are Sleeping Beauty transposons based on pITRSB that carry two genes: (***i***) an EFS-HygR gene and (**ii**) a TRE3G regulated gene designed to express codon optimized ORFs that encode WT CD63, CD63-YQRF, CD63-YQTI, and CD63-AEMV. All genes were synthesized *in vitro*, cloned into the appropriate, expression vectors, and then sequenced in their entirety to ensure the absence of unwanted mutations.

The plasmid used for knockout of the SDCBP gene, pJM1087, was based on pFF(*3*) and contains three genes. The first of these consists of a CMV promoter driving expression of a single long quadricistronic ORF that encodes (***i***) Cas9-3xNLS, (***ii***) a viral 2a peptide, (***iii***) EGFP, (***iv***) another viral 2a peptide, (***v***) the thymidine kinase (tk) from herpes simplex virus (HSV), (***vi***) another viral 2a peptide, and (***vii***) the puromycin resistance protein PuroR(*69*), with the ORF flanked by a pair of loxP sites. The second of these genes consists of the PolIII-transcribed H1 promoter diving expression of a Cas9 gRNA with the target sequence of 5’-ATAAACCTACTTCCATCGTG-3’, which is complementary to a sequence in the second coding exon of the SDCBP gene. The third of these genes consists of the PolIII-transcribed 7sk promoter diving expression of a Cas9 gRNA with the target sequence of 5’-GGTTTCTGGTGCACCACTTC-3’, which is complementary to a sequence in the third coding exon of the SDCBP gene.

Plasmids carrying genomic DNA (gDNA) amplification products from mutant cell lines were generated by extracting gDNA from single cell clones, amplifying small fragments of the genome surrounding the targeted site, using Taq polymerase. The resulting PCR fragments were checked for proper size, then inserted into a bacterial cloning vector by (A3600, Promega).

### Cell lines, transfections, and small molecules

293F were obtained from Thermo (A14528). SK-MEL-5 cells, Daudi cells, NIH3T3, an Hela-S cells were obtained from ATCC (HTB 70, CCL-213, CRL-1658, and CCL-2.2, respectively). The HEK293 Alix^-/-^ cell line was described previously(*3*), as was the 293F/CD63^-/-^ cell line (*50*). The 293F/AP2M1^-/-^ cell line, 293F/CD9^-/-/-^ cell line, and 293F/SDCBP^-/-^ cell line were generated in this report (see section below describing Cas9-mediated gene editing). Adherent cultures were grown in tissue culture plates in complete medium (DMEM, 10% fetal bovine serum, 1% penicillin/streptomycin) at 37°C, 90% humidity, and 5% CO_2_. For suspension cultures of 293F and 293F-derived cell lines, cells were grown in Freestyle medium (Thermo) in Erlenmeyer shaker flasks at 110 rpm, 37°C, 90% humidity, and 8% CO_2_. DNA transfections were performed using Lipofectamine 3000 according to the manufacturer’s instructions. Zeocin was used at 200 ug/mL. Puromycin was used at 3 ug/mL. Doxycycline was used at 10 ng/mL. Latrunculin A was used at 1 uM. Bafilomycin A was used at 100 nM.

To create the doxycycline-inducible Tet-on cell lines (i.e. FtetZ, 3TetZ, StetZ, HtetZ/Alix^-/-^, FtetZ/CD63^-/-^, and FtetZ/CD9^-/-/-^), the parental cell lines (293F, NIH3T3, Hela-S, HEK293/Alix^-/-^, 293F/CD63^-/-^, and 293F/CD9^-/-/-^, respectively) were transfected with pJM1463 using Lipofectamine 3000 (Thermo). Two days later, the transfected cell populations were placed in zeocin-containing media. The culture medium was changed every 3-4 days for 7-12 days to select for zeocin-resistant cells. The thousands of surviving single cell clones from each transfection were then pooled to generate a single polyclonal Tet-on derivative of each parental cell line.

To create the cell lines that express the syntenin proteins, FtetZ, 3TetZ, StetZ, and HtetZ/Alix^-/-^ cells were transfected with pYA128, YA129, and pYA130. Two days later the cells were placed in puromycin-containing media, followed by selection of puromycin-resistant clones and pooling of all clones to create polyclonal cell lines designed for the doxycycline-induced expression of syntenin proteins. To create the cell lines that express CD9 proteins, FtetZ/CD9^-/-/-^ cells were transfected with pCG606, pCG732, pCG733, pCG734, and pCG607. Two days later the cells were placed in hygromycin-containing media, followed by selection of hygromycin-resistant clones and pooling of all clones to create polyclonal cell lines designed for the doxycycline-induced expression of CD9 proteins. To create the cell lines that express CD63 proteins, FtetZ/CD63^-/-^ cells were transfected with pCG602, pMG9, pCG604, and pMG10. Two days later the cells were placed in hygromycin-containing media, followed by selection of hygromycin-resistant clones and pooling of all clones to create polyclonal cell lines designed for the doxycycline-induced expression of CD63 proteins.

### sEV preparation

For suspension cell cultures, cells were seeded into 30 ml of Freestyle medium at a density of 1 x 10^6 cells per ml and grown for 48-72 hours, with shaking. Culture media was collected and cells and cell debris were removed by centrifugation at 5,000 *g* at 4°C for 15 minutes and by passage of the resulting supernatant through a 200 nm pore size diameter filtration unit. To collect sEVs by size exclusion chromatography and filtration, the 200 nm filtrate was concentrated ∼100-fold by centrifugal flow filtration across a 100 kDa pore size diameter filter (Centricon-70, MilliporeSigma), followed by purification by size exclusion chromatography using qEV nano columns (Izon Sciences).

For adherent cell cultures, 6 million cells were seeded onto 150 mm dishes in 30 ml of complete medium, allowed to adhere to the plates overnight, then incubated for three days in complete medium. Culture media was collected and cells and cell debris were removed by centrifugation at 5,000 *g* at 4°C for 15 minutes and by passage of the resulting supernatant through a 200 nm pore size diameter filtration unit. The sEVs were then collected by differential centrifugation, supernatants were spun for 30 minutes at 10,000 x *g*, spun a second time for 30 minutes at 10,000 x *g*, then spun at 100,000 *g* for 2 hours, all at 4°C. The supernatant was discarded and the sEV pellet was resuspended for further analysis.

### Immunoblot

Cells were lysed in Laemmli/SDS-PAGE sample buffer lacking reducing agent. Samples were either maintained in reducing agent-free sample buffer or adjusted to 5% ß-mercaptoethanol, then heated to 100°C for 10 minutes, spun at 13,000 x g for 2 minutes to eliminate insoluble material, then separated by SDS-PAGE and processed for immunoblot as previously described using antibodies specific for human CD63 (NBP2-32830, NOVUS), mouse CD63 (NVG-2, Biolegend), CD9 (312102, Biolegend), CD81 (555675, BD Biosciences), Hsp90 (sc-13119, Santa Cruz Biotechnology), syntenin (PA5-76618, Thermo) AP2M1 (68196, Cell Signaling Technologies), Lamp1 (H4A3, Thermo), and Lamp2 (H4B4, Thermo). HRP-conjugated secondary antibodies were from Jackson ImmunoResearch.

### qRT-PCR

Total RNA was isolated using Quick-RNA Microprep Kit (Zymo Research). RNA was converted to single stranded cDNA by reverse-transcription using the High-Capacity RNA-to-cDNA Kit (Applied Biosystems). qPCR analysis was performed using SYBR Green master mix (Bio-Rad) and the CFX96 Real-Time PCR Detection System (Bio-Rad), with gene-specific primers for our codon optimized syntenin transgene (5′-GGCTCAAGTCTATTGATAATGGC-3′ and 5′-CCTTATCACTGGACCAACC-3′), our CD9 transgene (5′-GAAATGTATTAAATATCTTCTGTTCGGTTT-3′ and 5′-CCGGCACCGATAAGTATATAAAC-3′) and control primers for human 18S rRNA (5′-CGGCGACGACCCATTCGAAC-3′ and 5′-GAATCGAACCCTGATTCCCCGTC-3′). Data was analyzed with ΔΔCT Method.

### Flow cytometry and fluorescence activated cell sorting

Cells were released by trypsinization (TrypLE, Thermo) and cell clumps were removed using a cell-strainer (Falcon Cat#352235). Approximately 500,000 cells were then concentrated by a brief spin at 400 x *g* for 5 minutes and resuspended in 100 uL of 4°C FACS buffer (1% FBS in PBS) containing 2 uL of FITC-conjugated anti-CD63 (clone H5C6), 2 uL APC conjugated anti-CD9 (clone H19a), 2 uL PE conjugated anti-CD81 (clone 5A6), or 2 uL PerCP conjugated anti-Lamp1 (clone H4A3) and 2 uL PE conjugated anti-Lamp2 (clone H4B4), all from Biolegend, for 30 min with gentle mixing every 10 min. Cells were washed 3 times with 1 mL of 4°C FACS buffer, with cells recovered by 400 x *g* spin for 4 min at 4°C. After the final wash, cells were resuspended in FACS buffer with 0.5ug/ml DAPI, and analyzed using CytoFLEX S flow cytometer (Beckman Coulter). Flow cytometry histograms were generated using FlowJo (v10.8.1). For separation by FACS, labeled cells were sorted into single cells in a 96 well plate, on the basis of high or low cell surface labeling for CD63 or CD9.

### Cas9-medited gene editing

To create the 293F/AP2M1^-/-^ cell line, we transfected 293F cells with a mixture of Cas9 protein (A36498, Thermo) and an AP2M1-targeting single guide RNA (sgRNA; target sequence of 5’-ACGTTAAGCGGTCCAACATT-3’) using lipofectamine CRISPRMAX (Thermo). Cells were cultured for several days, then seeded into wells of into 96 well plates at one cell per well. Single cell clones (SCCs) were expanded, genomic DNA (gDNA) was extracted from each clone, and each gDNA was interrogated by PCR using AP2M1 gene-specific primers. PCR products were ligated into the pGEM®-T vector using a TA cloning kit (A3600, Promega), transformed into *E. coli*, and 8 or more clones from each ligation were sequenced in their entirety. The 293F/AP2M1^-/-^ cell line carried for further analysis carried one allele with a 198 bp deletion and one allele with = 1 codon deletion at a conserved position (Ile63) of the AP2M1 protein (***Fig. S5***).

A similar procedure was used to generate the 293F/CD9^-/-/-^ cell line. 293F cells were transfected with a mixture of Cas9 protein (A36498, Thermo) and a sgRNA guide RNA (target sequence of 5’-ATTCGCCATTGAAATAGCTG-3’) using lipofectamine CRISPRMAX (Thermo). Cells were cultured for several days, trypsinized, and seeded into wells of into 96 well plates at one cell per well. Single cell clones were expanded, genomic DNA (gDNA) was extracted from each clone, and each gDNA was interrogated by PCR using CD9 gene-specific primers. PCR products were ligated into the pGEM®-T vector using a TA cloning kit (A3600, Promega), transformed into *E. coli*, and 24 or more clones from each ligation were sequenced in their entirety. All gDNA amplification products from 293F/CD9^-/-/-^ cell lines carried up to three different sequences, indicating that 293F cells carry three CD9 alleles. The cell line used for further experimentation carried one CD9 allele with a 4 bp deletion and two CD9 alleles with the same 8bp deletion. (***Fig. S8***).

The 293F/SDCBP^-/-^ cell line was created by transfecting 293F cells with the plasmid pJM1087 then selecting for puromycin-resistant cell clones. After 7 days in selection, surviving cells were pooled, with EGFP-positive cells separated by FACS into individual wells of a 96 well plate. The emergent SCCs were expanded and 10 were interrogated by immunoblot using antibodies specific for syntenin, revealing that all 10 lacked detectable level of syntenin protein. We then extracted gDNA from these clones and interrogated each gDNA by PCR using primers flanking both gRNA target sites in the SDCBP gene. The 293F/SDCBP^-/-^ cell line selected carried deletions between the two target sites in coding exons 2 and 3 on both of its SDCBP alleles rendering both functionally null. To delete the Cas9-EGFP-HSVtk-PuroR ORF, this cell line was transfected with a Cre recombinase expression vector, grown for 10 days in CM lacking antibiotics, then seeded at various densities into CM containing ganciclovir to eliminate any HSVtk-expressing cells. After growth and selection for two weeks in ganciclovir-containing medium, the resulting cell clones were pooled and examined by flow cytometry, revealing that all cells in the population were EGFP-negative. Furthermore, exposure of these cells to puromycin confirmed that all of the cells in the ganciclovir-resistant population had also reverted to puromycin sensitivity, and were therefore likely no longer expressing Cas9.

### Endocytosis assay and confocal fluorescence microscopy

Cells were grown overnight on poly-D-lysine-coated coverglasses in CM, then transferred to pre-chilled 4°C CM. Cells were washed in 4°C PBS, then incubated for 30 minutes at 4°C with pre-chilled 4°C PBS (200 uL) containing 4 uL FITC-conjugated anti-CD63 (clone H5C6, Biolegend) and 4 uL APC conjugated anti-CD9 (clone H19a, Biolegend). Excess antibody was removed by two washes with 4°C PBS. Cells were then fixed immediately or transferred to CM at 37°C and incubated for 30 minutes at 37°C. Fixation was with 3.7% formaldehyde in PBS for 20 minutes. Cells were then incubated with DAPI to stain the nucleus, and then examined by confocal fluorescence microscopy and imaged to assess the subcellular distribution of plasma membrane-labeled CD63 and CD9. Confocal fluorescence microscopy was performed using a Zeiss LSM880 microscope with gallium-arsenide phosphide (GaAsP) detectors and a 63x/1.4na Plan-Apochromat objective. Images were assembled into figures using ImageJ and Adobe Illustrator.

### SPIR-IFM analysis

Exosomes were purified by filtration and size exclusion chromatography, then further purified by affinity capture on anti-CD81 or anti-CD9 antibodies that had been immobilized on SPIR imaging chips (Unchained Labs). Bound exosomes were then stained using CF-647 conjugate of a mouse anti-human CD63 monoclonal antibody (Unchained Labs) and imaged using an Exoview R200 imaging platform (Unchained Labs).

### qSMLM analysis

25 mm diameter coverslips #1.5H (Thermo Fischer Scientific, Cat# NC9560650; Waltham, MA, USA) were functionalized with N-hydroxysuccinimide (NHS) groups, followed by covalent attachment of monoclonal antibodies that bind to epitopes in the ectodomain of human CD81 and human CD9. FtetZ cells and FtetZ::syntenin cells were grown in Freestyle media containing doxycycline, followed by collection of their exosomes by concentrating filtration and size exclusion chromatography. The resulting exosome preparations were diluted in PBS containing 0.025% Tween 20 (1:50) to a final volume of 80 uL and placed on the surface of antibody-coated coverslips at room temperature overnight in a humidified chamber. Coverslips were then washed with PBS containing 0.025% Tween 20 and EVs were labeled with a cocktail of AF647-labeled antibodies specific for human CD63 (Novus Biologicals Cat. No. NBP2-42225 Centennial, CO, USA) and CF568-labeled antibodies specific for human CD9 (Bio<u>L</u>egend, Cat. No. 312102, San Diego, CA, USA) and human CD81 (BioLegend, Cat. No. 349502, San Diego, CA, USA). All antibodies were fluorescently labeled as described previously at a molar ratio of ∼1 (*70*)). Samples were fixed and stored as described previously(*71, 72*).

For imaging, coverslips were placed in Attofluor cell chambers (Thermo Fisher Scientific, Cat. No. A7816) loaded with direct stochastic optical reconstruction microscopy (dSTORM) imaging buffer(*73*). N-STORM super-resolution microscope (Nikon Instruments; Melville, NY, USA) was used for SMLM imaging using 561 nm and 640 nm lasers, respectively, using microscope components described previously(*74*). Images were acquired using NIS-Elements software (Nikon Instruments). SMLM images were processed using N-STORM Offline Analysis Module of the NIS-Elements software to localize peaks as described before(*71*). The localization data were analyzed with the Nanometrix software (version 1.0.4.61; Nanometrix Ltd, Oxford, UK) using a density-based spatial clustering of applications with noise (DBSCAN) algorithm. Prior to cluster analysis, the detected 640 nm and 561 nm channel localizations were aligned using the Uniform Dual Channel Alignment tool of Nanometrix. The DBSCAN-based cluster identification was performed at a neighbor search radius of 30 nm and minimum points per cluster of 30 as analysis conditions. Post-processing of the detected cluster data, including concatenation and filtering, was performed using Matlab (version R2022a; MathWorks; Natick, MA, USA). Clusters were identified as EVs considering the following constraints. In the case of the 647 nm channel, the minimum and maximum number of localizations per cluster was set to 30 and 3000, as well as the minimum and maximum diameter was set to 20 nm and 400 nm, respectively. In the case of the 561 nm channel, the minimum and maximum number of localizations per cluster was set to 40 and 3200, as well as the minimum and maximum diameter was set to 30 nm and 400 nm, respectively. Colocalized (CD63+, CD81/CD9+) EVs were identified as overlapping clusters detected in the 640 nm and 561 nm channel. The number of tetraspanin molecules per exosome was calculated using an average of 15 (647 nm channel) and 16 (561 nm channel) localizations per single fluorescent tetraspanin antibody, as described before(*75*). Following EV identification, the EV count per region of interest (ROI), CD63 molecule count per EV, and EV diameter were further analyzed. To account for the difference in EV concentration across the samples, we normalized the number of detected CD63+, CD81/CD9+ and CD63-, CD81/CD9+ EVs by the total number of CD81/CD9+ EVs. Statistical significances in the normalized EV count per ROI data were determined using Brown-Forsythe and Welch ANOVA test. Statistical significance in the CD63 molecule count per EV and diameter data was assessed performing two-tailed Welch’s t-test after logarithmic transformation. Statistical analysis and graph generation were performed in GraphPad Prism (version 9.5.1; GraphPad, San Diego, CA, USA).

### Analysis of RNAseq data

The web tool ‘AnalyzeR’ was used to analysis RNA expression data of CD63 and SDCBP in >230,00 distinct samples that had been examined from normal and malignant human tissues, as previously described(*76*).

## Supporting information

Supplemental material for Ai et al. revision

## Acknowledgments

The authors thank James Morrell for his outstanding technical assistance and to the many colleagues who provided feedback over the course of this project, especially Drs. Xandra Breakefield, Michael Caterina, Chulhee Choi, Saumya Das, Samir El-Andaloussi, Dhanu Gupta, Tijana Jovanovic-Talisman, Michiel Pegtel, Shang Jui Tsai, and Antje Zickler.

## Author contributions

Conceptualization: YA, CG, MG-C, SJT, TJ-T, SJG

Methodology: YA, CG, MC-G, ‘AS, TJ-T, SJG

Investigation: YA, CG, OS, SR, YD, SJG

Visualization: YA, CG, MG-C, AS, SJG

Funding acquisition: TJ-T, SJG

Project administration: TJ-T, SJG

Supervision: TJ-T, RL, SJG

Writing – original draft: SJG

Writing – review & editing: YA, CG, MG-C, TJ-T, SJG

## Data and materials availability

All data are available in the main text or the supplementary materials.

