## Supplemental material for Ai et al. revision for "Syntenin and CD63 Promote Exosome Biogenesis from the Plasma Membrane by Blocking Cargo Endocytosis"

Yiwei Ai *et al*.

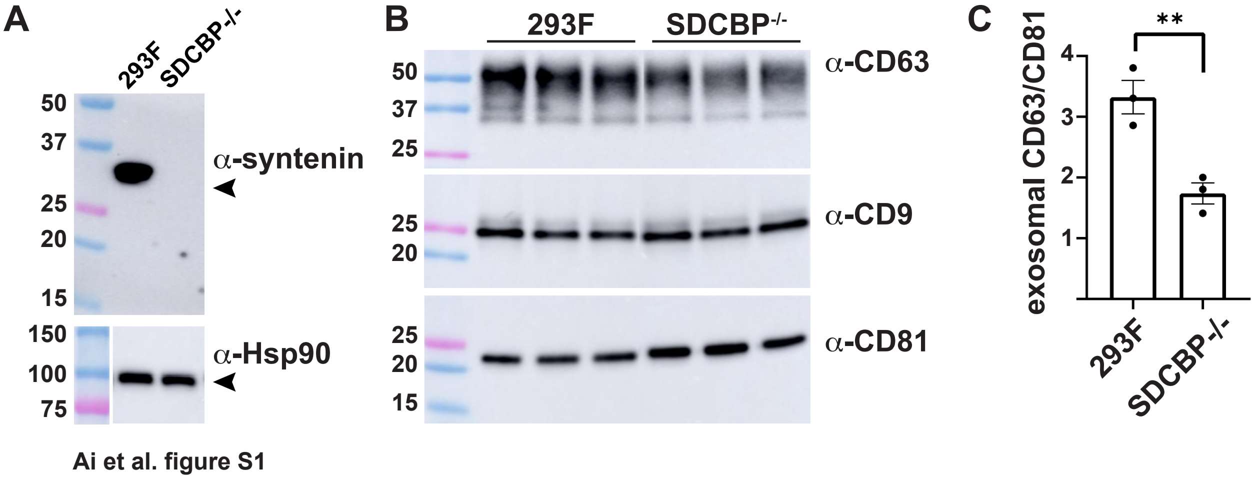


**Figure S1. Knockout of the SDCBP gene in 293F cells results in an ~40% decrease in the vesicular secretion of CD63**. (**A**) Immunoblots of 293F cells and a 293F SDCBP$^{-/-}$ cell line probed with antibodies specific for syntenin and for the cytoplasm protein Hsp90. The SDCBP$^{-/-}$ cell line has large indel mutations on both allele #1 and allele #2 of the SDCBP gene, both of which delete the 3' half of coding exon 2 and the 5' half of coding exon 3. MW markers are in kDa. (**B**) Immunoblot analysis of exosomes collected from triplicate cultures of 293F cells and the F/SDCBP$^{-/-}$ cell line, probed with antibodies specific for CD63, CD9, and CD81. 293F cells and 293F/SDCBP$^{-/-}$ cells were grown in triplicate in Freestyle medium, with shaking, for three days. MW markers are in kDa. (**C**) Bar graph showing the relative exosomal ratio of CD63:CD8, with bar height representing the average, and error bars showing the standard error of the mean. Individual data points are shown, and the asterisks (**) refer to a Student's test $p$ value <0.01. Data are from three independent trials.

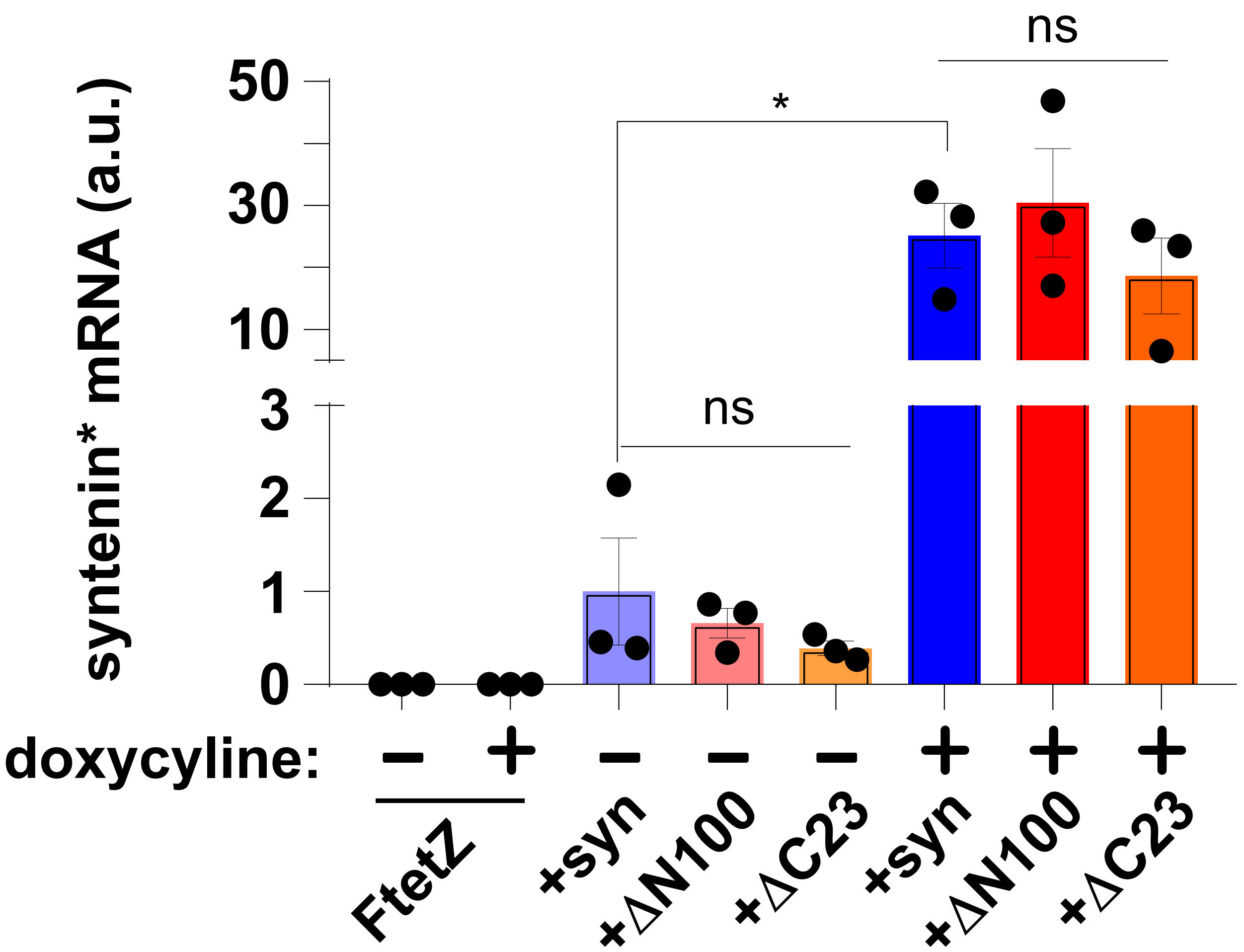


**Figure S2. Abundance of transgene-encoded syntenin mRNAs in uninduced and induced cells.** Bar graph of transgene-encoded syntenin mRNA levels (syntenin*) in control FtetZ cells and FtetZ cells carrying doxycycline-inducible transgenes encoding syntenin (syn), ΔN100 syntenin (DN100) or synteninΔC23 (DC23). Bar height represents the average, error bars showing the standard error of the mean, and * represents a *p* value <0.05. Individual data points are shown. Data are from three independent trials.

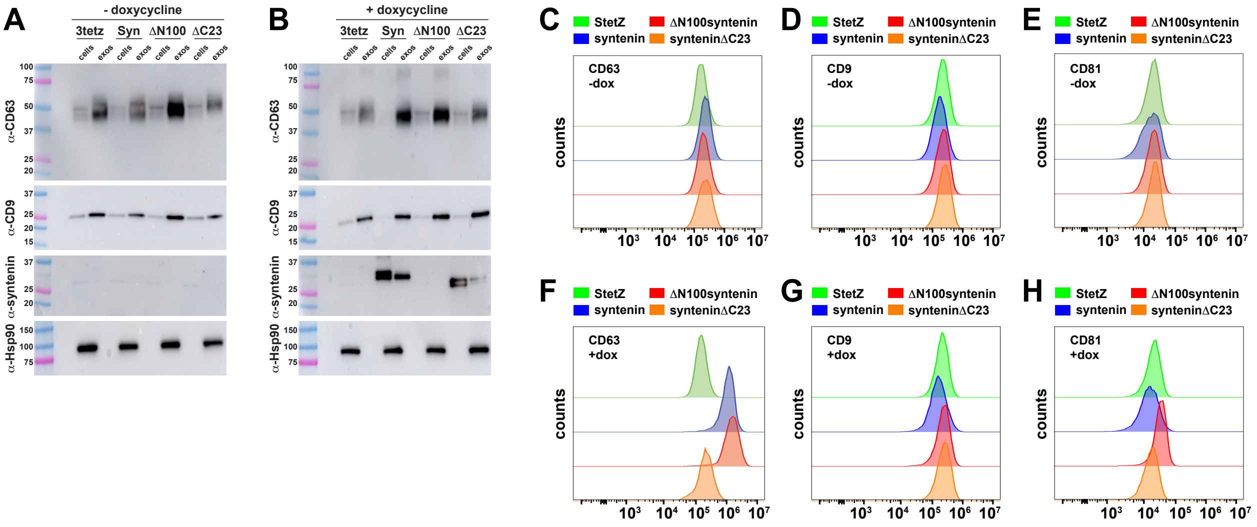


**Figure S3. Syntenin induces the exosomal secretion and plasma membrane accumulation of CD63.** (**A, B**) Immunoblot analysis of cell and exosome lysates collected from a Tet-On derivative of mouse NIH3T3 cells, 3TetZ, and also from 3TetZ cells carrying TRE3G-regulated transgenes encoding (Syn) WT human syntenin, (ΔN100) ΔN100syntenin, or (ΔC23) synteninΔC23. Cells had been grown for 3 days in the (A) absence or (B) presence of doxycycline, and probed using antibodies specific for CD63, CD9, syntenin, and Hsp90. Data are from three independent trials. (**C-H**) Flow cytometry histograms of (green) Tet-On Hela-S cells (StetZ) and StetZ cells carrying TRE3G-regulated transgenes encoding (blue) WT human syntenin, (red) ΔN100syntenin, or (orange) synteninΔC23, that had been grown in the (C-E) absence or (F-H) presence of doxycycline, then chilled to 4ºC and stained using (C, F) FITC-labeled antibodies specific for CD63, (D, G) PE-labeled antibodies specific for CD81, or (E, H) APC-labeled antibodies specific for CD81. Data are from three independent trials.

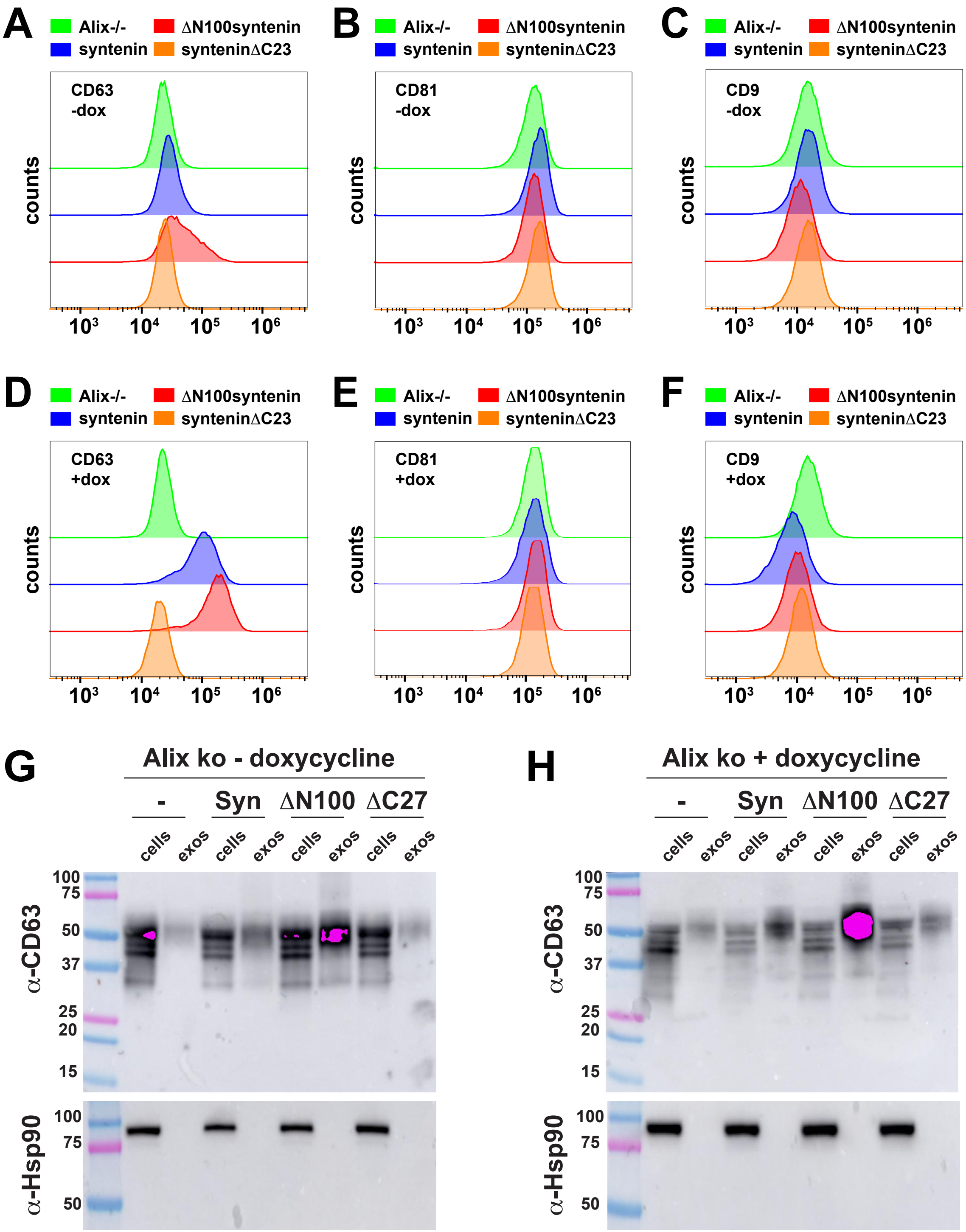


**Fig. S4. Syntenin induces the biogenesis of CD63 exosomes by an Alix-independent mechanism**. (**A-F**) Flow cytometry histograms of (green) HtetZ/Alix$^{-/-}$ cells and HtetZ/Alix$^{-/-}$ cells carrying TRE3G-regulated transgenes encoding (blue) WT human syntenin, (red) ΔN100syntenin, or (orange) synteninΔC23, that had been grown in the (**A-C**) absence or (**D-F**) presence of doxycycline, then chilled to 4ºC and stained using (A, D) FITC-labeled antibodies specific for CD63, (B, E) PE-labeled antibodies specific for CD81, or (C, F) APC-labeled antibodies specific for CD9. (**G, H**) Immunoblot analysis of cell and exosome lysates collected from (-) HtetZ/Alix$^{-/-}$ cells and HtetZ/Alix$^{-/-}$ cells carrying TRE3G-regulated transgenes encoding (Syn) WT human syntenin, (ΔN100) ΔN100syntenin, or (ΔC23) synteninΔC23, that had been grown in the (G) absence or (H) presence of doxycycline, and probed using antibodies specific for CD63 or Hsp90. Data are from three independent trials.

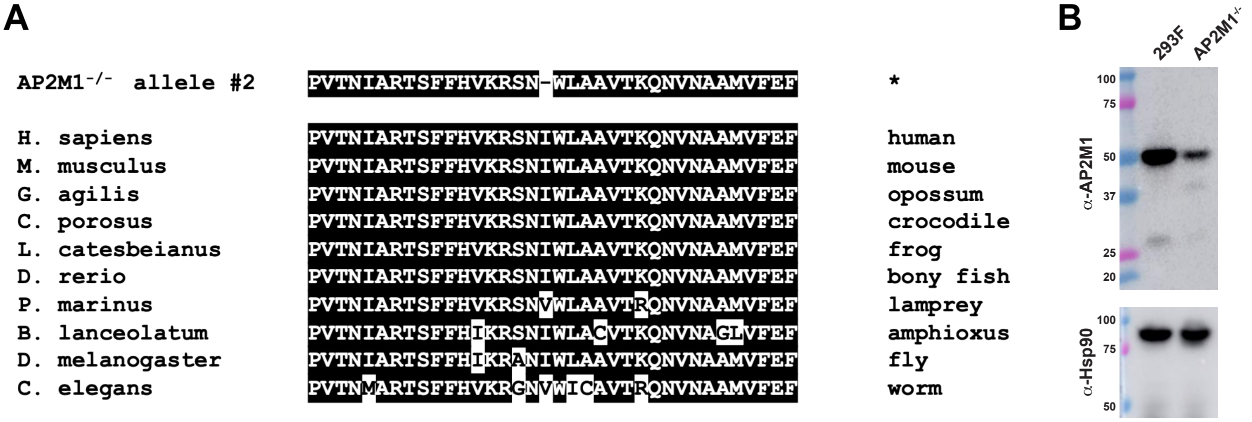


**Figure S5. Characterization of an AP2M1$^{-/-}$ derivative of 293F cells**. (**A**) Amino acid alignment showing (top line) the 1 amino acid deletion (I63Δ) encoded by allele #2 of the AP-2 *mu* protein and (lower lines) the same region of the AP-2 mu2 protein encoded by other metazoans. Please note the highly conserved spacing of sequences flanking I63, as well as the presence of Ile or Val at this position in each of these metazoan homologs. (**B**) Immunoblot of cell lysates from 293F cells and the 293F AP2M1$^{-/-}$ cell line, probed with antibodies specific for AP2M1 and Hsp90. Protein molecular weight markers are listed by size in kDa. Note that the amount of AP2M1 protein present in the AP2M1$^{-/-}$ cell lines is greatly reduced but not absent, which is expected for a cell line that carries a null mutation on allele #1 of the AP2M1 gene (a 198 bp deletion that eliminates the 5' half of exon 3, the 3' end of intron 2, and precludes proper splicing of the AP2M1 transcript) and a 1 codon deletion on allele #2 that encodes a mutant AP2M1 protein of nearly normal length. Data are from two independent trials.

**A**

```
           V  I  F  A  I  E  I  A  A  A  I  W  G  Y  S  H  K  D  E  V  I  K
```

WT GT**GATATTCGCCATTGAAATAGCTG**CGGCCATCTGGGGATATTCCCACAAGGATGAGGTGATTAAG

Allele #1 GTGATATTCGCCATTGAA....CTGCGGCCATCTGGGGATATTCCCACAAGGATGAGG**TGA**TTAAG

Allele #2 GTGATATTCGCCATTGAAATAG........ATCTGGGGATATTCCCACAAGGA**TGA**GGTGATTAAG

Allele #3 GTGATATTCGCCATTGAAATAG........ATCTGGGGATATTCCCACAAGGA**TGA**GGTGATTAAG

**B**

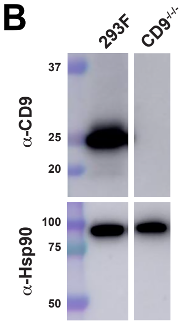


**Figure S6. Genetic and biochemical validation of a 293F CD9$^{-/-}$ cell line**. (**A**) Amino acid and DNA sequences of (top line) the WT human CD9 gene in the vicinity of the CD9-targeting gRNA target site (bold) and PAM site (underlined) that was used to mutate the CD9 gene, positioned above the DNA sequences of the three CD9 alleles in the 293F/CD9$^{-/-}$ cell line (293F cells carry three alleles of the CD9 gene). The sequences of the three mutated CD9 alleles were determined by amplifying a genomic DNA amplicon flanking the gRNA target site, followed by cloning the amplification product into a bacterial cloning vector, a then sequencing 25 separate clones of the amplicon, of which 6 clones carried the 4 bp deletion and 19 clones carried the 8 bp deletion. (**B**) Immunoblot of cell lysates extracted from the parental 293F cell lines and the 293F/CD9$^{-/-}$ cell lines, probed with antibodies specific for CD9 and Hsp90. Protein molecular weight markers are listed by size in kDa. Data are from two independent trials.

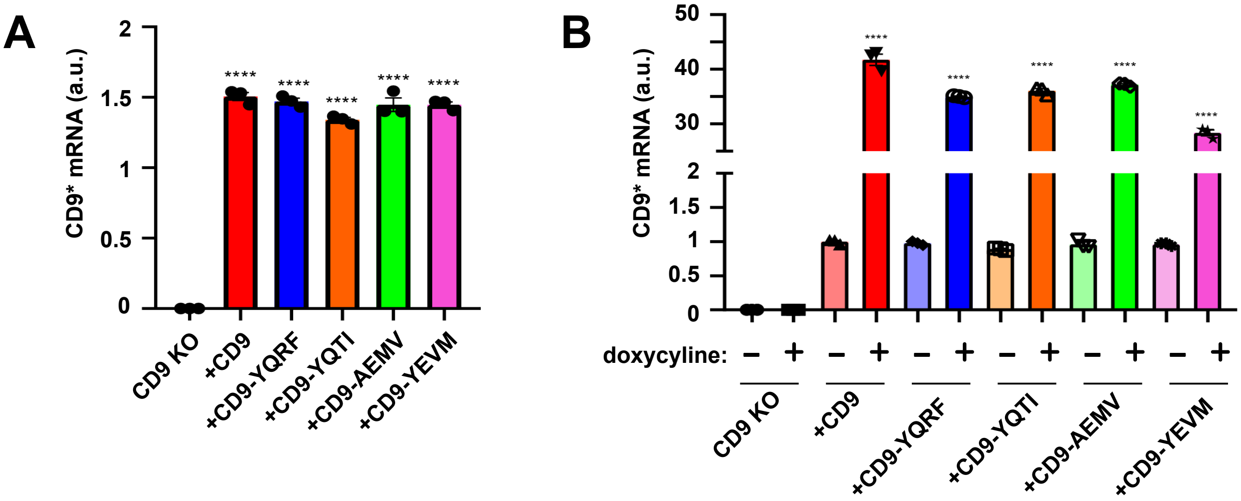


**Figure S7. Relative abundance of transgene-encoded CD9 mRNAs in uninduced and induced cell lines**. (**A**) Bar graph of CD9 transgene-encoded mRNA abundance, as determined by qRT-PCR, in uninduced FtetZ/CD9$^{-/-/-}$ cells and uninduced FtetZ/CD9$^{-/-/-}$ cells carrying TRE3G-regulated, codon-optimized CD9 genes encoding (red) WT CD9, (blue) CD9-YQRF, (orange) CD9-YQTI, (green) CD9-AEMV, or (purple) CD9-YEVM. (**B**) Bar graph of CD9 transgene-encoded mRNA abundance, as determined by qRT-PCR, in uninduced and doxycycline-induced FtetZ/CD9$^{-/-/-}$ cells alone or in FtetZ/CD9$^{-/-/-}$ cells carrying TRE3G-regulated, codon-optimized CD9 genes encoding (red) WT CD9, (blue) CD9-YQRF, (orange) CD9-YQTI, (green) CD9-AEMV, or (purple) CD9-YEVM. Data are from three independent trials.

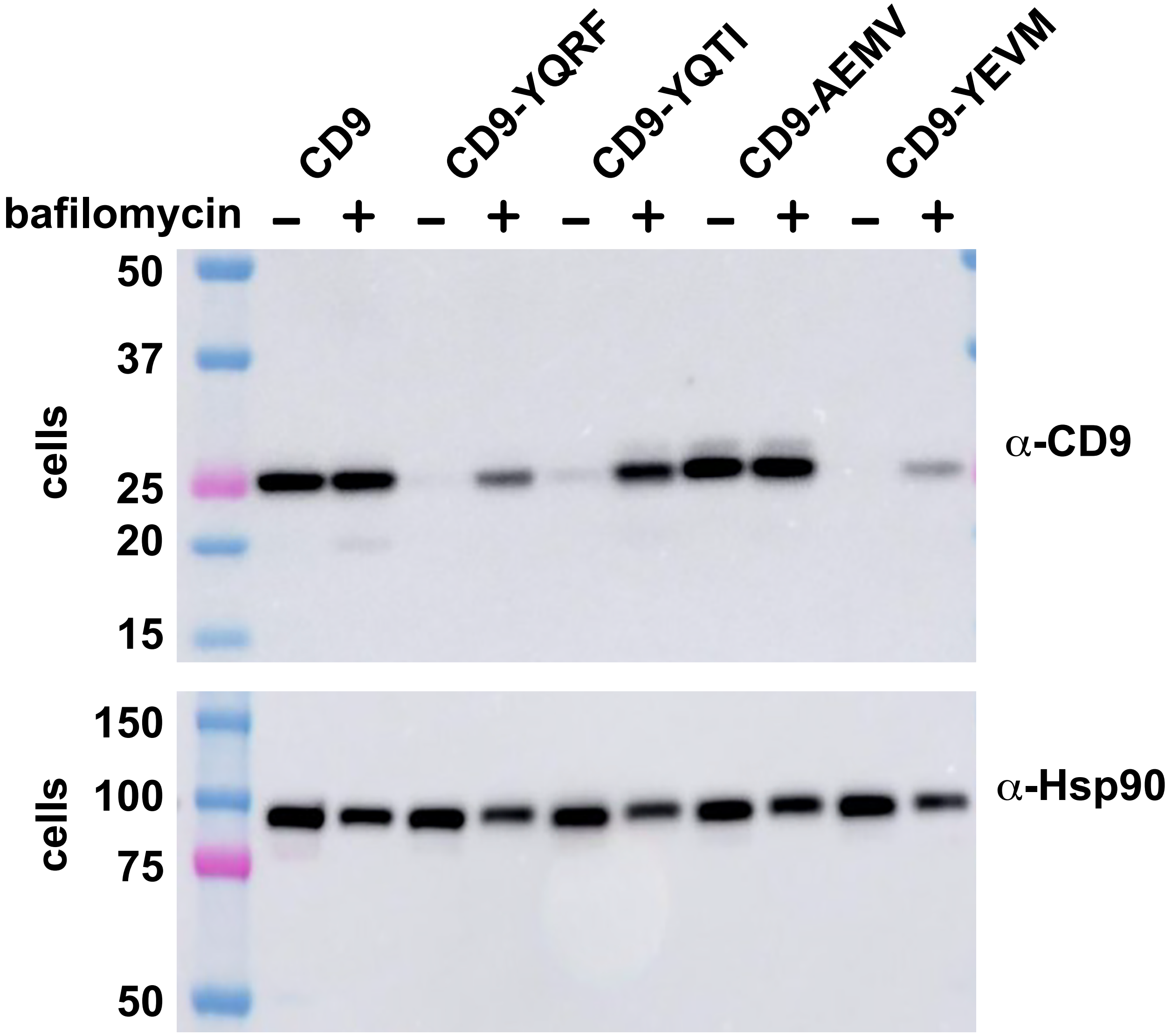


**Figure S8. Effect of bafilomycin on the abundance of transgene-encoded CD9 proteins**. FtetZ/CD9$^{-/-/-}$ cells carrying TRE3G-regulated, codon-optimized CD9 genes encoding WT CD9, CD9-YQRF, CD9-YQTI, CD9-AEMV, or CD9-YEVM were grown in normal media in the absence or presence of bafilomycin, then lysed and processed for immunoblot using antibodies specific for CD9 or Hsp90. Protein molecular weight markers are listed by size in kDa. Data are from three independent trials.

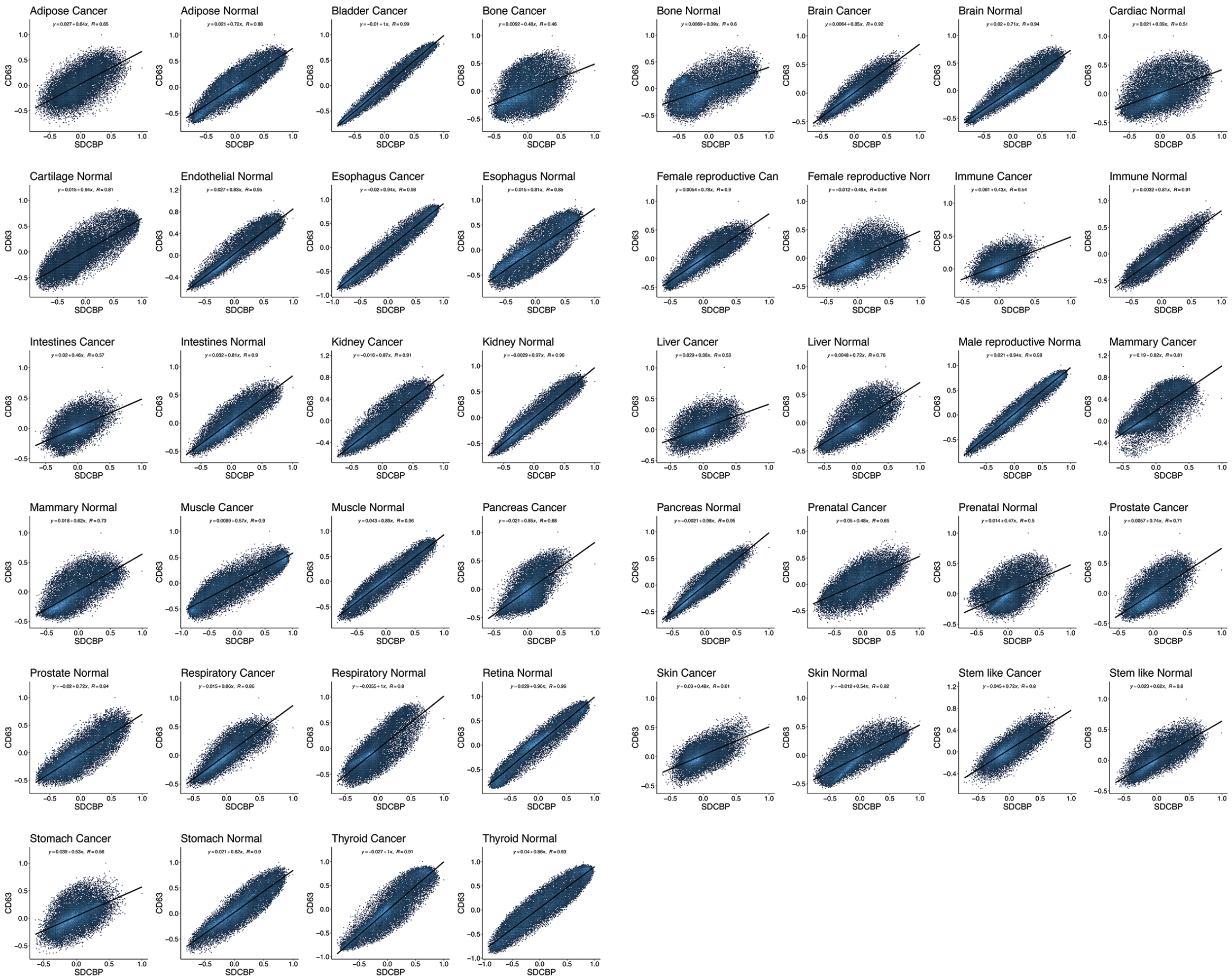

**Figure S9. Syntenin and CD63 mRNA levels display a strong positive correlation across >200,000 RNA samples collected from 44 sets of normal tissue samples and tumor samples**. Scatter plots of log2 mRNA abundance for (y axis) CD63 and (x-axis) syntenin (SDCBP) across diverse RNAseq databases. Values refer to the y-axis intercept, the slope as a function of x, and the Pearson correlation coefficient (*R*).

**Table S1. CD63 transcript levels in selected human cell types.** Data from proteinatlas.org.

| Cell type | CD63 mRNA abundance in nTPM |
|---|---|
| Syncytiotrophoblasts | 3398 |
| Basal respiratory cells | 3151 |
| Exocrine glandular cells | 2810 |
| **SK-MEL-5** | **2373** |
| Melanocytes | 2259 |
| Leydig cells | 1798 |
| Granulocytes | 1770 |
| Fibroblasts | 1444 |
| Cardiomyocytes | 1276 |
| Smooth muscle cells | 1206 |
| Macrophages | 1164 |
| Cholangiocytes | 878 |
| Adipocytes | 800 |
| Endothelial cells | 795 |
| Breast myoepithelial cells | 613 |
| Schwann cells | 564 |
| Granulosa cells | 514 |
| Hepatocytes | 434 |
| Monocytes | 372 |
| Keratinocytes | 341 |
| T-cells | 306 |
| **HEK293** | **249** |
| B-cells | 185 |
| Dendritic cells | 132 |
| Astrocytes | 27 |
| **Daudi** | **87** |
| Oligodendrocytes | 25 |
| Microglia | 11 |
| Neurons | 9 |

**Table S2. The cargo-based analysis of exosome biogenesis presented here supports the shared, stochastic model of exosome biogenesis, not the endocytosis/exocytosis model of exosome biogenesis**

| empirical observation | prediction of the shared, stochastic model | prediction of the endocytosis/exocytosis model |
|---|---|---|
| Syntenin expression induced the exosomal secretion of CD63 by blocking its endocytosis | **YES** | NO |
| Knockout of AP2M1 induced the exosomal secretion of CD63 | **YES** | NO |
| Latrunculin induced the exosomal secretion of CD63 | **YES** | NO |
| Syntenin-induced the exosomal secretion of CD63 independently of Alix | **YES** | NO |
| Syntenin is not required for the exosomal secretion of CD81 or CD9 | **YES** | NO |
| Appending endocytosis signals to CD9 decreased its exosomal secretion | **YES** | NO |
| YxxΦ forms of CD9 induced the PM accumulation and exosomal secretion of CD63 | **YES** | NO |
| High level expression of CD63 inhibited its own endocytosis | **YES** | NO |
| High level expression of CD63 inhibited AP-2-mediated endocytosis in general | **YES** | NO |
| Removing CD63's YxxΦ motif induced its plasma membrane accumulation and exosomal secretion | **YES** | NO |
| Removing CD63's YxxΦ motif reduced its ability to inhibit AP-2 endocytosis | **YES** | NO |
| Naturally variation in CD63 expression regulates the cell surface accumulation of Lamp1 and Lamp2 | **YES** | NO |
